# Spatially-Organized Adaptive Immune Niches in Granulomas Dictate Local Defense and Correlate with Disease Severity in Tuberculosis Patients

**DOI:** 10.64898/2026.11.16.766905

**Authors:** Furong Qi, Yingyin Cao, Qianting Yang, Hao Wang, Ruipeng Huang, Kun Qiao, Mingfeng Liao, Zhiqiang Cheng, Wei Zhang, Tantan Ren, Geyang Luo, Juanjuan Zhao, Diana Boraschi, Lázaro Moreira Marques Neto, Yuedong Yang, Bo Yan, Zheng Zhang

**Author notes:** These authors contributed equally (Furong Qi, Yingyin Cao, Qianting Yang, Hao Wang, Ruipeng Huang, Kun Qiao). Correspondence (Z.Z.); (F.Q.); (B.Y.).

## Abstract

The cellular composition and spatial organization of immune responses within tuberculosis (TB) granulomas have been increasingly characterized, yet how these spatially organized immune cell populations coordinate local adaptive immunity remains largely unknown. Here, we integrated single-cell multi-omics, spatial transcriptomics, and immune receptor profiling to construct a comprehensive spatial atlas of human TB granulomas. We identified two concentric adaptive immune niches surrounding the central necrotic core. The outer niche comprised granuloma-associated tertiary lymphoid structures (TLSs) that supported T_FH_-B cell interactions and exhibited functional germinal center-like features associated with antigen-driven B-cell maturation toward FCRL5⁺ antibody-producing cells. The inner niche contained antigen-experienced and *Mtb*-reactive effector memory CD8⁺ T cells (T_EM_), which were recruited from distal lung regions by *LGALS9*^high^ chemokine-expressing macrophages and activated to undergo local clonal expansion. In *lgals9l1* deficient zebrafish, macrophage depletion was accompanied by marked reductions in T– and B-cell enrichment within granulomas and increased bacterial loads, supporting a role for *LGALS9*^high^ macrophages in coordinating local adaptive immunity against TB infection. Finally, AI-based prediction of spatial gene expression from clinical hematoxylin and eosin-stained specimens revealed that inferred TLS abundance increased with TB disease severity. These findings identify spatially interconnected adaptive immune niches that organize protective immune responses within human TB granulomas, therefore providing a foundation for understanding granuloma adaptive immunity and developing host-directed therapies and vaccines for TB (https://tbomics.smart-nbc.org.cn/).

## Introduction

Tuberculosis (TB), caused by the intracellular pathogen *Mycobacterium tuberculosis* (*Mtb*), remains one of the leading causes of infectious-disease mortality worldwide, accounting for more than 1.5 million deaths annually^1^. Following infection, the local immune landscape is initially shaped by host-pathogen interactions within heterogenous cell populations^2–4^. Especially, the tissue-resident alveolar macrophages (AMs) provide a nutrient-rich intracellular niche that can support *Mtb* persistence and impair their phagolysosomal maturation^4–6^, whereas inflammatory monocyte-derived macrophages (MMs) are recruited to sites of infection and establish a more restrictive antimicrobial microenvironment through the production of pro-inflammatory cytokines and reactive oxygen^7^. Importantly, these early myeloid responses initiate an inflammatory cascade that recruits diverse adaptive immune cells into the lung parenchyma^8–10^. Their coordinated accumulation ultimately gives rise to granulomas, the pathological hallmark of TB; however, the comprehensive spatial organization of these immune cells within granulomas has not been well documented especially at single cell resolution.

Granulomas are dynamic immunological microenvironments with paradoxical functions^8^. They physically compartmentalize infected cells and constrain bacterial dissemination while simultaneously providing permissive niches that support *Mtb* persistence, metabolic adaptation, and immune evasion. Determining how granuloma architecture is organized and remodeled to balance bacterial containment against progressive tissue pathology remains a central challenge in TB immunology^2^. Histological studies have characterized granulomas as structured immune cell aggregates comprising a macrophage-rich core surrounded by lymphocytes and fibroblasts^10–12^. Recently, several single-cell RNA sequencing (scRNA-seq) studies in zebrafish^13–16^, mice^17–21^ and non-human primates (NHPs)^14,17,22–30^ have provided important insights into granuloma biology, but these animal models cannot fully mirror human granuloma and the immune responses and disease dynamics in human TB granuloma remains largely uncharacterized. Emerging spatial-omics studies have delineated distinct cellular communities and the layered architecture within human TB granulomas with limited markers^3,31–34^. Nevertheless, the complex immune microenvironment of these lesions, particularly their adaptive immune features, remains poorly resolved without paired scRNA-seq and immune repertoire profiling.

Innate effector mechanisms, including those mediated by macrophages, are insufficient for durable control of *Mtb*. Effective containment additionally requires spatially organized adaptive immune niches that coordinate T– and B-cell responses with infected myeloid cells. Several studies have shown that the timely induction, spatial recruitment, and functional execution of *Mtb*-specific adaptive immunity can influence the transition between bacterial containment and progressive disease^5,35^. For decades, the CD4⁺ T cell-IFN-γ-macrophage axis has served as a canonical framework for protective immunity against TB^29,36,37^. *Mtb* infection can elicit long-lasting protection against subsequent exposure depending on CD4^+^ T cells^25,38–40^. In particular, Th1, Th17 and stem cell-like memory T cells and cytotoxic T cells are important for bacterial control^24^. Although this paradigm remains central to our understanding of host defense against *Mtb*, accumulating evidence has highlighted additional CD8^+^ T cell populations and effector programs that contribute to antimycobacterial immunity, particularly during early protection^27^. *Mtb*-specific CD8⁺ T cells can directly target infected cells through perforin-, granzyme-, and granulysin-dependent cytotoxic mechanisms, with also production of inflammatory cytokines and chemokines that contribute to local immune coordination^24,27,41^. Together, these studies establish *Mtb*-specific CD8⁺ T cells as an important determinant of granuloma-level bacterial control, but they leave unresolved a fundamental spatial question: how are protective CD8^+^ T cells selectively recruited and positioned within granulomas to access their cellular targets?

Accumulating evidence also implicates a key role of B cells in *Mtb* immunity, although their contributions appear to be highly context– and site-dependent^34,42–47^. In mice, B cell deficiency can alter pulmonary lesion development, inflammatory cell recruitment, bacterial dissemination and disease severity. However, these findings have been inconsistent, potentially owing to differences in experimental variables, including inoculum dose and *Mtb* strains^45,46,48^. In nonhuman primates, B cells can regulate granulomatous responses^47^, and antigen-specific B cells may contribute more to anti-*Mtb* defense than previously appreciated through their colocalization and cooperation with local T follicular helper (T_FH_)-like cells^45,46^. Human granuloma B cells produce *Mtb*-reactive antibodies to facilitate phagocytosis^42^. Emerging evidence further suggests that effective immune control is associated with ectopic tertiary lymphoid structures (TLSs) surrounding granulomas^34,43–45^. These findings suggest that B cells are integral components of the adaptive immune response to *Mtb*. However, how B cell niches are spatially organized within human granulomas and how their interactions with T_FH_ cells and other immune populations contribute to local host defense remain poorly defined.

Here, we integrated single-cell multi-omics, spatial transcriptomics, and immune-repertoire profiling to construct a comprehensive spatial atlas of human TB granulomas. We resolved two interconnected adaptive immune niches surrounding the central necrotic core: an inner antigen-reactive T_EM_ cell-macrophage niche and an outer TLS compartment characterized by T_FH_-B cell interactions and germinal center like features. We further found that *LGALS9*^high^ chemokine-expressing macrophages play a key role in the organization of these adaptive immune niches. We further trained an hematoxylin and eosin (HE)-based artificial intelligence (AI) model to predict spatial gene expression in clinical species and found that inferred TLS abundance positively correlates with TB severity. An interactive online resource (https://tbomics.smart-nbc.org.cn/) is developed to explore the cellular and spatial features of human TB granulomas. These findings establish a spatial framework linking innate and adaptive immunity and provide a foundation for understanding granuloma immunity and developing of host-directed therapies and vaccines.

## Results

### Single-cell analysis reveals prominent enrichment of adaptive immune cells in human pulmonary TB granulomas

To dissect the immune cell compositions and track changes during human TB granuloma formation, twenty-four biopsy samples were collected from nine patients with active pulmonary TB during surgery (**Table S1**). The samples were analyzed using single-cell RNA sequencing (scRNA-seq), single-cell B cell receptor sequencing (scBCR-seq), and single-cell T cell receptor sequencing (scTCR-seq) (**Figure 1A**). For each patient, active lesion tissues (TBL, n = 9), tissues from the lesion boundary or adjacent areas (TBA, n = 6), and tissues located distal to the lesions (TBD, n = 9) were collected. In addition, tissues from six healthy lung samples from deceased donors (HD) were included as controls. In total, 264,787 immune cells passed quality control and were integrated into a joint map (**Figure 1B**).

**Figure 1.**
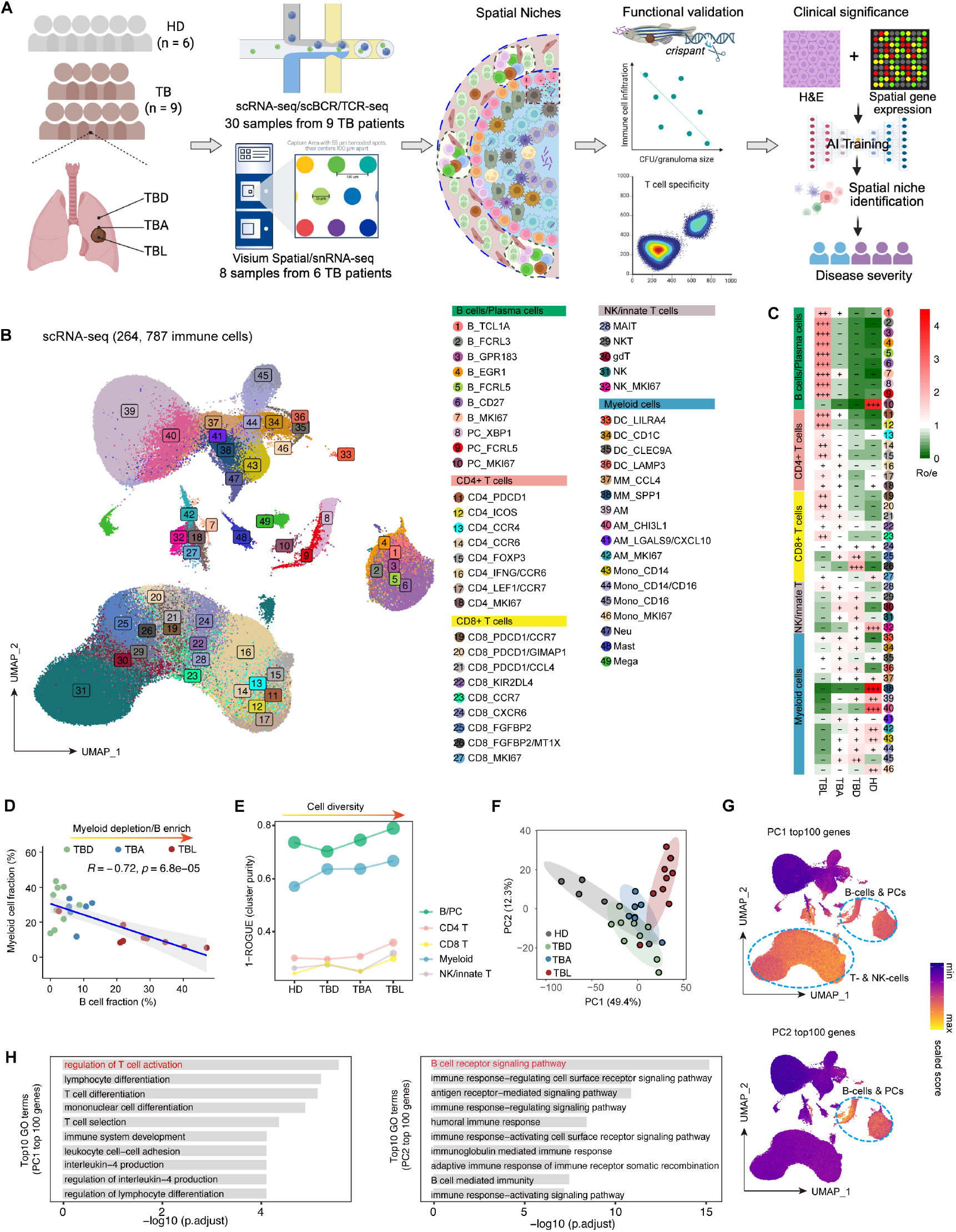
Single-cell analyses reveal prominent adaptive immune alterations in granulomatous lesions. (A) Schematic overview of the study. Spatial transcriptomics, snRNA-seq, immunofluorescence staining and AI-based spatial gene-expression prediction were performed on FFPE samples, whereas scRNA-seq and scBCR/TCR-seq were performed on surgically resected granuloma tissues. (B) Immune-cell landscape identified by scRNA-seq. A total of 49 immune-cell subtypes were resolved. (C) Heatmap showing the enrichment of cell subtypes based on observed-to-expected ratios (Ro/e). Ro/e < 1 (-); 1 < Ro/e <= 1.5 (+); 1.5 < Ro/e <= 2 (++); Ro/e > 2 (+++). (D) Correlations between the cell fractions of B cells and myeloid cells. (E) Cluster purity of five major cell types estimated using ROGUE. (F) Principal component analysis of all samples based on the mean expression of highly variable genes. PC1, principal component 1. PC2, principal component 2. (G) Module score projection on the UMAP for the top 100 genes contributing to PC1 (top) and PC2 (bottom). (H) The 10 most significantly enriched Gene Ontology terms (p.adjust < 0.05) identified from the top100 genes contributing to PC1 (left panel) and PC2 (right panel).

We identified 49 cell types shared across patients (**Figure 1B and S1A**, **Table S2**). Among these, the proportions of B cells, plasma cells (PCs), and T cells significantly increased in TBL compared to the other groups, while macrophages and monocytes showed the most significant decrease in TBL (**Figure 1C** and **S1B**). Notably, a transition from a macrophage-dominated to a B cell-dominated microenvironment was observed from TBD to TBL (**Figure S1C**). Consistent with this observation, the macrophage and B cell fractions were negatively correlated across lung tissues, indicating that B cells were preferentially enriched in TBL where granulomas commonly occur (**Figure 1D**). More importantly, the cell heterogeneity as measured by the ROGUE metric was also significantly increased in TBL across all the cell populations (**Figure 1E**), indicating increased cellular heterogeneity and complexity within human TB granulomas.

The transcriptional profiles further distinguished TBL from TBD and TBA in a principal component analysis (PCA) based on the average expression of the 2,000 most variable genes (**Figure 1F**). TBL was separated from TBA and TBD along principal component (PC) 2. To investigate the immune features underlying this separation, we explored the immune characteristics of the top 100 genes associated with PC1 and PC2. The average gene expression levels of the top 100 genes on PC1 were predominantly expressed by T and B cells, whereas the top 100 genes on PC2 were uniquely expressed by B cells and PCs (**Figure 1G**). Functional annotation using Gene Ontology (GO) analysis showed that these genes were enriched in processes related to T-cell and B-cell activation (**Figure 1H**). Collectively, these results indicate that human granuloma lesions exhibit distinctive adaptive immune features that differentiate them from surrounding and distal lung tissues.

### Macrophage-CD8 T_EM_ and T_FH_-B cell niches are recurrent features of TB granulomas across patients

To identify immune cell populations that co-occurred in TBL areas, we applied BIRCH clustering to their frequency profiles. This analysis resolved three groups of co-occurring immune cell populations (**Figure 2A**). Cluster 1 is primarily composed of peripheral helper T cells (CD4_ICOS), T_FH_ cells (CD4_PDCD1), and all identified B cell populations. Cluster 2 mainly includes effector memory CD8^+^ T cells (T_EM_, CD8_PDCD1) expressing the inhibitory or exhausted receptors such as *TIGIT* and *PDCD1*, T helper 1 cells (T_H_1, CD4_IFNG/CCR6), regulatory T cells (Treg, CD4_FOXP3), FCRL5^high^ plasma cells (PC_FCRL5), and specially chemokine-expressing alveolar macrophages (AM_LGALS9/CXCL10). Cluster 3 comprised monocyte derived macrophages (MM_CCL4, MM_SPP1), monocytes (Mono_CD14, Mono_CD14/CD16, Mono_CD16), natural killer cells (NK), dendritic cells (DC_CD1C, DC_LAMP3) and effector CD8^+^ T cells (CD8_FGFBP2).

**Figure 2.**
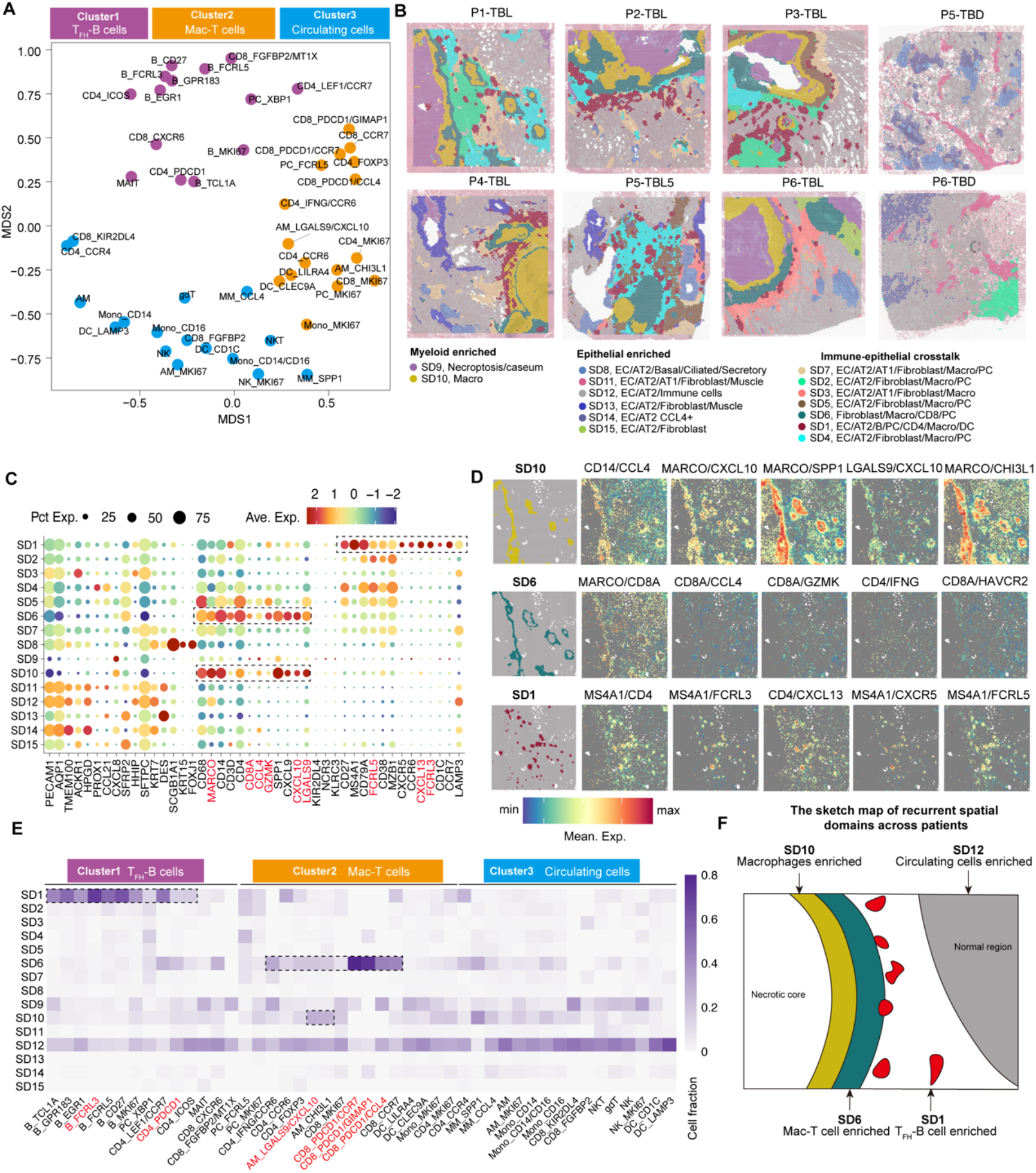
Adaptive immune-cell niches in granulomatous lesions. (A) Clustering of cell frequencies across 49 immune-cell subtypes. Three immune-cell clusters were identified by BIRCH clustering in TBL samples. Cluster 1 and 2 were predominantly tissue-resident cell populations, whereas cluster 3 consisted primarily of circulating cell populations. (B) Spatial domains inferred using MNMST across six granuloma lesions (TBL) and two distal normal tissue (TBD) sections. (C) Dot plot showing indicated gene expression patterns across spatial domains. Pct.Exp., percent of cells expressing the gene. Ave.Exp., average gene expression. (D) Spatial heatmaps showing the mean expression of two-marker combinations in P1-TBL. (E) Relative cell-type fractions in each spatial domain based on cell type decomposition analysis. (F) Schematic representation of spatial domains shared across patients.

To resolve the spatial organization of these populations, we performed spatial transcriptomics sequencing on six granuloma lesions and two distal control regions (**Table S1**, **Figure 1A**). Using a multi-layer network model^49^, we identified 15 spatial domains (SDs) across these 8 samples (**Figure 2B**). SD1, SD6, and SD10 were uniquely present within TBL region across all six patients compared to TBD area, indicating that they represent recurrent features of granuloma organization. Several marker genes helped define these spatial domains (**Table S3**). In brief, *MARCO*, *CXCL10* and *LGALS9* defined chemokine-expressing alveolar macrophages (hereafter referred to as *LGALS9*^high^ macrophages); *MS4A1*, *FCRL5*, *FCRL3* defined B cells; and *CXCL13* and *CD4* defined T_FH_ cells; *CD8A*, *CCL4* and *GZMK* defined CD8^+^ T_EM_ cells (**Figure 2C**). These signatures, together with the average expression of corresponding marker panels in shared spatial spots, distinguished macrophage-enriched domains (SD6 and SD10), a B cell enriched domain (SD1), and a domain containing both macrophages and T_EM_ cells (SD6) (**Figure 2D** and **S2**). These spatial expression patterns suggested that some domains corresponded to the co-occurring immune cell populations identified by scRNA-seq. SD1 was consistent with Cluster 1. Both SD6 and SD10 were associated with *LGALS9*^high^ alveolar macrophages, but only SD6 showed prominent T_EM_-associated expression, linking this domain to Cluster 2. By contrast, except *SPP1*, markers associated with Cluster 3 were distributed across multiple domains, with no corresponding discrete spatial compartment.

To further assess these associations, we defined a gene panel representing each cell population identified from our scRNA-seq data (**Table S4**). For each cell population, we calculated the relative enrichment fraction based on the proportions of spots containing these cell markers across the 15 SDs (**Methods**). B cell populations are most enriched in SD1, while T_FH_ cells are enriched in both SD1 and SD6 (**Figure 2E**). Macrophage subpopulations, such as AM_*LGALS9/CXCL10* and AM_*CHI3L1*, are primarily located within SD6 and SD10, whereas T_EM_ cells predominantly associate with SD6. Cluster 3 cell populations (monocytes, and some NK cells, DCs and effector CD8^+^ T cells) were distributed across multiple domains, including those in distal control tissue (**Figure 2E**), and were therefore excluded from subsequent niche-focused analyses.

Given that individual spatial transcriptomic spots can capture multiple cells, we further assessed cellular composition using single-nucleus RNA sequencing (snRNA-seq) of adjacent tissue sections from the same six donors (**Figure 1A**). This analysis yielded 44,448 nuclei and resolved immune populations alongside epithelial, stromal, and vascular populations (**Figure S3A**). The latter included basal, alveolar type I (AT1), alveolar type II (AT2), ciliated, and secretory cells; fibroblasts and muscle cells; and lymphatic, capillary, aerocyte, and venous endothelial populations (**Figure S3B**). We used this reference to deconvolve the cellular composition of spatial transcriptomic spots^50^ (**Figure S3C**). Consistent with the marker-based analysis, *FCRL3*^high^ B cells and T_FH_ cells (CD4_PDCD1) were enriched in SD1. Chemokine-expressing AM (Marco_CXCL12) and T_EM_ cells (CD8_CCL4) were enriched within SD6 and SD10, whereas epithelial populations characteristic of distal normal lung tissue were enriched in SD12.

Together, these data provide an unprecedented view of the spatial organization of human TB granulomas at single-cell resolution. We defined two adaptive immune niches surrounding the necrotic core: a T_FH_-B cell niche in SD1 and a T_EM_-macrophage niche in SD6 (**Figure 2F**). Our spatial analysis also highlights *LGALS9*^high^ macrophages as an underappreciated cell population in granuloma organization.

### The T_FH_-B cell interaction niche exhibits TLS features

Previous studies have shown that *Mtb* infection can promote the formation of B-cell follicles and TLSs within tubercular granulomas^34,43,51,52^. However, the cellular and molecular organization of TLSs within granulomas remains incompletely defined. In particular, it is unclear whether they contain functionally active germinal centers.

Compared with HDs, T_FH_ cells (CD4_PDCD1) in TBL samples exhibited stronger inferred interactions, particularly with B_FCRL5 cells (**Figure 3A**). Among the inferred ligand-receptor interactions, the CXCL13-CXCR5 axis was the dominant signaling pathway mediating communication between T_FH_ and B-cell populations at both the single-cell level (**Figure 3B**) and within the SD1 spatial domain (**Figure 3C**). Immunofluorescence staining further confirmed the close spatial relationship between T_FH_ cells and B cells, in which T_FH_ was identified by CD4 and CXCL13, whereas B cells were identified by CD20 and CXCR5 (**Figure 3D**). Given the established role of CXCL13 in recruiting CXCR5-expressing lymphocytes and organizing lymphoid follicles, enhanced CXCL13-CXCR5 signaling suggests the presence of an active T_FH_-associated TLS-like niche within granulomas^45,53,54^.

**Figure 3.**
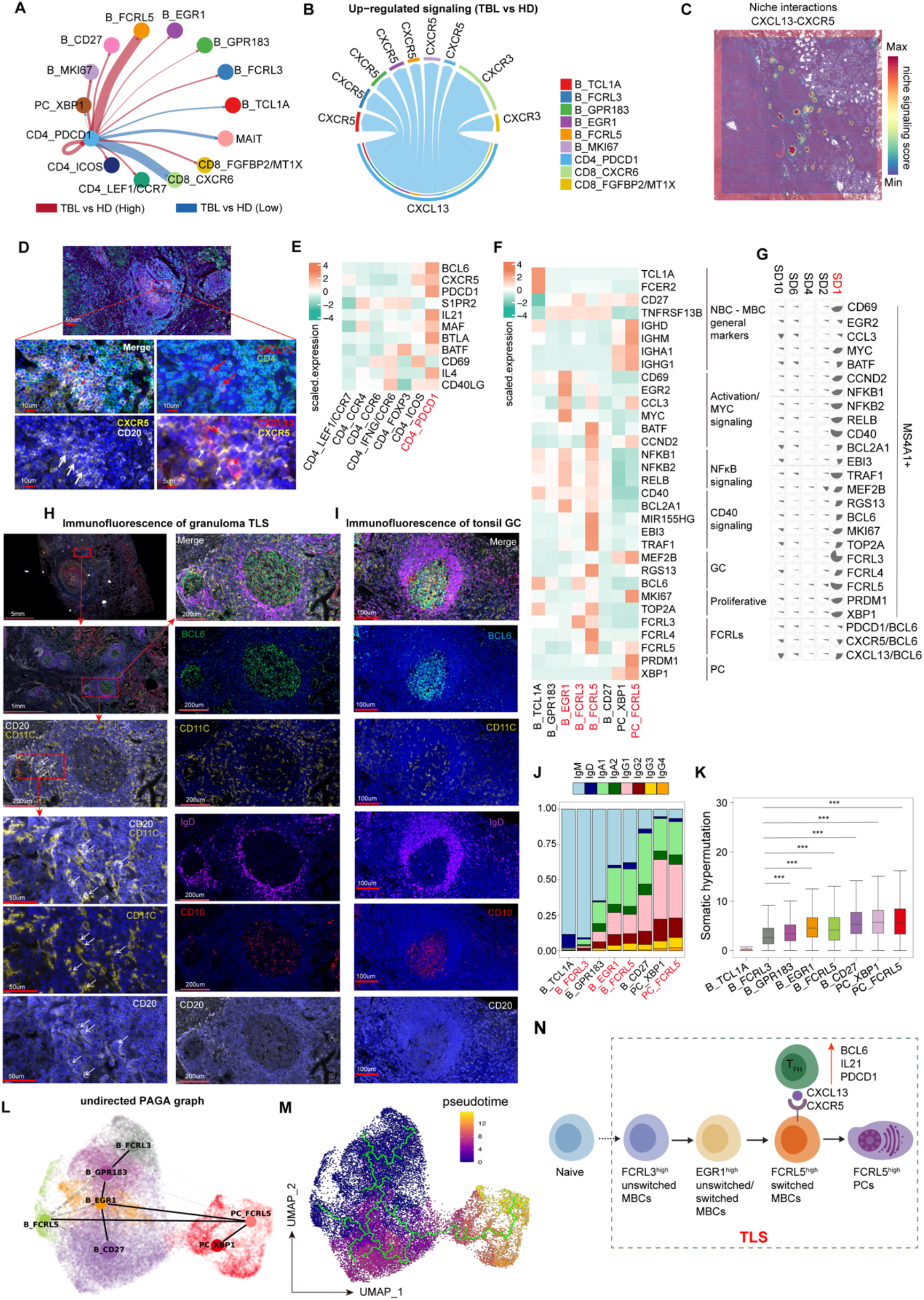
The T_FH_-B cell interplay niche displays functional TLS features. (A-B) CellChat v2-inferred interaction strengths and upregulated ligand-receptor pairs using T_FH_ cells as the source population. Red and blue lines indicate interactions that were significantly stronger and weaker, respectively, in TBL than in HD samples. (C) Spatial interaction scores for CXCL13-CXCR5 signaling inferred using NICHES. (D) Immunofluorescence staining showing CXCL13^+^ T_FH_ cells (red arrows) and CXCR5^+^ B cells (white arrows) in granulomas. (E) Mean expression of GC reaction-related genes across CD4^+^ T cell subsets. (F) Average expression of indicated functional genes across B cell subsets. (G) Relative cell-type fractions inferred by cell decomposition analysis in selected spatial domains, where the sector size indicates cell fraction. (H-I) Immunofluorescence staining showing TLS structures (H), atypical memory B cells (CD11C^+^ B cells highlighted), and tonsil GC architecture (I). (J) Relative antibody isotype frequencies across B-cell subsets. (K) Somatic hypermutation rates in heavy-chain V genes from TB patients compared with germline sequences across B-cell states. Statistical significance was calculated using Mann-Whitney U Test (\*\*\**P* < 0.001) using B_FCRL3 as control. (L) PAGA graph showing undirected connectivity among B-cell subtypes. (M) Single-cell trajectories inferred using Monocle3, with B_FCRL3 cells designated as the root population. (N) Schematic representation of the inferred B-cell differentiation pathway within TLSs.

Among CD4^+^ T-cell subsets, T_FH_ cells displayed the highest expression of canonical GC-associated genes, including *BCL6*, *IL21*, *MAF*, *BTLA*, *CXCL13* (**Figure 3E**). They also had the lowest TCR diversity (**Figure S4A**), and showed pronounced clonal expansion (**Figures S4B** and **S4C**), consistent with ongoing antigen-driven selection^55^. To characterize the functional states of B-cell populations, we performed gene set-based analyses (**Figure S4D**, **Table S5**). B_TCL1A represented naïve B cells, whereas B_GPR183 corresponded to early activated mature B cells. B_EGR1 displayed an activated memory B-cell program characterized by enrichment of MYC and NF-κB signaling, consistent with the role of *EGR1* as an immediate-early regulator downstream of BCR activation^56^. B_FCRL5 exhibited features of atypical B cells (ABCs), which have been described in chronic infection, autoimmunity, and cancer^57,58^. They also expressed a pre-GC transcriptional program marked by *CCND2*, and *MIR155HG*, together with increased MYC activity (**Figure 3F, Figure S4D**). B_FCRL3 cells preferentially expressed follicular-associated genes, including the GC master regulator *BCL6*, together with several inhibitory BCR signaling molecules. Compared with PC_XBP1, PC_FCRL5 retained early GC-associated features, such as *MEF2B*, while also expressing programs associated with PC differentiation and antibody production, including *PRDM1*. These GC-associated populations (B_FCRL3, B_FCRL5, and PC_FCRL5) were significantly expanded in TBL regions (**Figures S4E** and **S4F**). We next examined whether these GC-associated transcriptional programs exhibited spatial organization. SD1 showed the strongest enrichment of GC signatures (**Figure S4G**). Spatial feature analyses demonstrated co-localization of *MS4A1* with GC-associated genes within SD1 (**Figures 3G** and **S4H**). T_FH_-associated markers and *BCL6* were also predominantly localized to SD1, supporting that this domain represents a T_FH_-B cell interaction niche with GC-like properties. Immunofluorescence staining further support for the presence of TLSs within granulomas by demonstrating the co-localization of CD20, BCL6, and CD10 (**Figure 3H**). We also observed a marked accumulation of ABCs within TLSs, identified by co-expression of CD20 and CD11c (**Figure 3H**). The granuloma-associated TLSs are very similar to GC structure in tonsil when staining with classical markers (**Figure 3I**). Notably, we found that TLSs exhibited a thicker IgD^+^ mantle zone and a substantially greater accumulation of perifollicular PCs than tonsil GCs.

To further determine whether granuloma-associated TLSs contain bona fide GCs, we analyzed two Visium HD datasets of normal human tonsils generated by 10x Genomics. Spatial clustering identified 14 anatomical regions, including follicular regions (clusters 0, 4, 8, 9, 13), interfollicular regions (clusters 2, 6 10) and epithelial regions (clusters 1, 3, 5, 7, 11, 12) (**Figure S5A**). Among the follicular regions, cluster 4 exhibited a dark-zone program characterized by low *CD83* and high *BCL6* expression, whereas cluster 0 displayed a light-zone program characterized by high *CD83 and low BCL6* expression. Cluster 8 corresponded to PC-committed light-zone B cells (**Figures S5B** and **S5C**). Clusters 9 and 13 contained myeloid and T-cell populations located within GCs. Projection of granuloma-derived B-cell populations onto these reference states revealed a continuum of GC-like transcriptional programs. B_FCRL3 most closely resembled dark-zone GC B cells, characterized by high *BCL6 and CXCR4* expression. B_FCRL5 exhibited both dark-zone (*CIDA*^high^) and light-zone (*LMO2*^high^) features, including expression of genes associated with GC re-entry (**Figure S5D**). B_EGR1 predominantly aligned with light-zone programs (**Figure S5D**). Spatial deconvolution further showed that tonsillar dark– and light-zone reference clusters (cluster 4 and 0, respectively) mapped primarily to SD1 (**Figure S5E**).

### T_FH_-B cell TLS niches exhibit functional germinal center-like features and support B-cell maturation toward FCRL5⁺ plasma cells

B cell follicles provide an anatomical niche for antigen-driven B cell selection, affinity maturation, and somatic hypermutation (SHM). Among the GC-like B cell populations identified in TB granulomas, B_FCRL3, B_EGR1, and B_FCRL5 exhibited substantially lower BCR diversity than other memory B-cell subsets (**Figure S6A**), consistent with ongoing antigen-driven clonal selection. Analysis of immunoglobulin isotypes revealed a progressive decline in IgM accompanied and an increase in IgG from B_FCRL3 to B_FCRL5, indicating a transition from unswitched to class-switched B cells (**Figure 3J**). PCs were almost exclusively class-switched and exhibited the highest SHM frequencies (**Figure 3K**), consistent with extensive affinity maturation. Higher SHM frequencies within GCs have been associated with increased BCR affinity and preferential differentiation toward plasmablasts rather than memory B cells^59,60^. Consistent with a bias toward memory-cell maintenance, B_FCRL3 exhibited the lowest SHM frequency among memory B-cell populations (**Figure 3K**). This interpretation was further supported by its transcriptional profiling. B_FCRL3 preferentially expressed *CD24* (**Figure S6B**), which has been reported to restrain plasmablast differentiation^61^, together with FCRL3, a receptor implicated in promoting B cell activation while limiting PC differentiation^62^. B_EGR1 expressed *IRF8*, a negative regulator of PC differentiation^63^. In contrast, B_FCRL5 showed robust expression of PC commitment genes, including *PRDM1*, *FAS*, and *IRF4*, consistent with progression toward the PC lineage^64^.

The spatial restriction of B_FCRL3 to TLSs, together with its transcriptional resemblance to dark-zone GC B cells, promoted us to hypothesize a sequential differentiation program in which B_FCRL3 gives rise to B_EGR1 and B_FCRL5, ultimately generating PC_FCRL5. Partition-based Graph Abstraction (PAGA) analysis^65^ supported a putative developmental trajectory from B_FCRL3 to B_EGR1, B_FCRL5, and ultimately PC_FCRL5 (**Figure 3L**). This trajectory was independently supported by Monocle3 pseudotime analyses^66^ (**Figure 3M**). Clonal tracking demonstrated extensive BCR sharing among these four populations (**Figure S6C**). In addition, reconstructed lineage trees based on CDR3 evolution were consistent with transitions among B_FCRL3, B_EGR1, and B_FCRL5, with PC_FCRL5 consistently occupying terminal branches (**Figures S6D** and **S6E**).

Collectively, these findings reveal that granuloma-associated TLSs provide organized GC-like niches that support B-cell maturation from FCRL3⁺ cells toward FCRL5⁺ antibody-producing PCs (**Figure 3N**). These findings highlight the potential GC-like function of granuloma-associated TLSs and suggest a mechanism for generating local humoral immunity during *Mtb* infection.

### Macrophage-T_EM_ interaction niches are associated with recruitment and activation of *Mtb*-reactive CD8^+^ T cells

We next investigated how macrophage-T_EM_ interactions shape local adaptive CD8⁺ T-cell immunity. Alveolar macrophages are important host cells for *Mtb* and contribute to bacterial control and granuloma organization. Our scRNA-seq data revealed substantial heterogeneity among myeloid cells, including monocytes, macrophages and dendritic cells (**Figure S7A)**. Among the macrophage populations, *LGALS9*^high^ AMs (AM_LGALS9/CXCL10) and *CCL4*^+^ MMs (MM_CCL4) progressively increased in frequency from HD to TBL samples (**Figure S7B**). Previous studies have identified *CCL4*^+^ MMs as a monocyte-derived inflammatory macrophage population with similar characteristics across different species^6,16,27,41^. Here, *LGALS9*^high^ AMs displayed a tissue-resident activation program (**Figure S7C**). Based on our curated macrophage gene signatures (**Table S5**), *LGALS9*^high^ AMs showed the strongest inflammatory and M1-like signatures, both at the single-cell level and within the corresponding spatial domains, SD6 and SD10 (**Figure S7C** and **S7D**). In addition to expressing inflammatory mediators such as *IL6*, *TNF*, and *IL1A*, this population was enriched for pathogen-recognition genes, including *TLR2*, *TLR4*, and *LGALS9* (**Figure S7E**), consistent with a potential role in sensing and responding to *Mtb* infection. Genes associated with epithelioid macrophage differentiation, including *MMP9* and *CD274*, were also most highly expressed in *LGALS9*^high^ AMs (**Figures S7F** and **S7G**), suggesting that this population may combine inflammatory activity with features associated with granuloma organization^67^. Consistent with recent reports, galectin-9, encoded by *LGALS9*, has been implicated in recognizing *Mtb*-induced phagosome damage, binding mycobacterial arabinogalactan, and promoting autophagy-mediated bacterial killing^68^.

The spatial co-occurrence of *LGALS9*^high^ macrophages and T_EM_ cells within SD6 prompted us to investigate whether these cell populations form a functionally relevant immune niche specially in response to *Mtb* infection. Cell-cell communication analysis revealed stronger inferred interactions between *LGALS9*^high^ macrophages (AM_LGALS9/CXCL10) and T_EM_ cells (CD8_PDCD1) in TBL than in HD samples (**Figure 4A**). At the single-cell level, the CXCL9/10/11-CXCR3 axis was among the dominant predicted signaling pathways connecting these populations (**Figure 4B**). The spatial analysis also supported a similar role for the LGALS9-TNFRSF9 axis within the macrophage-T_EM_ niche (**Figure 4C**). Immunofluorescence further confirmed the spatial co-localization of CXCL10 expressing CD68⁺ macrophages and TIGIT-expressing CD8^+^ T_EM_ cells around granulomatous lesions (**Figures 4D and S8A**). Because CXCL10 can recruit activated CXCR3⁺ T cells to inflamed tissues^69^, these findings are consistent with chemokine-producing macrophages in the accumulation of CD8⁺ T cells within granulomas.

**Figure 4.**
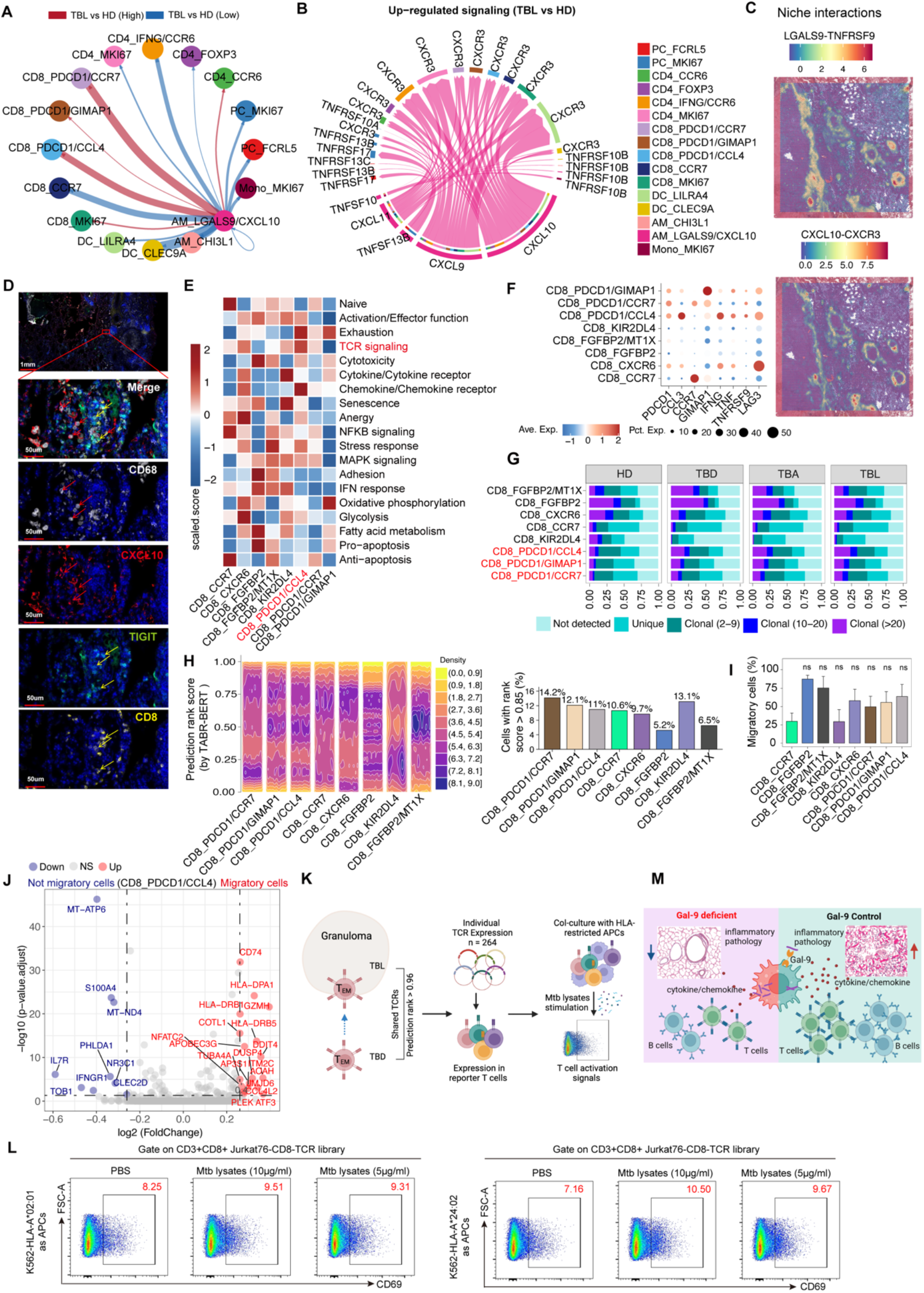
Macrophage-T_EM_ interplay niche promotes activation of *Mtb*-reactive T_EM_ cells. (A-B) CellChat v2-inferred cell-cell interaction strength using Macro_LGALS9/CXCL10 cells as the source population. Red and blue lines indicate interactions that were significantly stronger and weaker, respectively, in TBL than in HD samples. (C) Spatial interaction scores for CXCL10-CXCR3 and LGALS9-TNFRSF9 signaling inferred using NICHES. (D) Immunofluorescence staining showing CXCL10^+^ macrophages and CD8^+^TIGIT^+^ T_EM_ cells in granulomas. Yellow arrows indicate T_EM_ cells. Red arrows indicate CXCL10^+^ macrophages. (E) Heatmap showing gene-set-based phenotypic scores across CD8^+^ T cell subtypes. (F) Dot plot showing the expression of exhaustion-associated genes (*PDCD1* and *LAG3*), and effector-function-associated genes (*IFNG* and *TNF*). Pct.Exp., percent of cells expressing the gene. Ave.Exp., average gene expression. (G) Stacked bar plot showing clonal-state distributions across CD8^+^ T cell subtypes. (H) Density plot of antigen-specificity rank scores inferred using TABR-BERT (left panel) and the fraction of cells with rank scores > 0.85 (right panel). Statistical significance was calculated using Mann-Whitney U Test (ns, not significant) using CD8_CCR7 as control. (I) Fraction of migratory cells across CD8^+^ T cell subsets. (J) Volcano plot showing DEGs between migratory and non-migratory T_EM_ cells. The significant up and down regulated genes (P < 0.05) are indicated by red and blue. (K) Experimental workflow used to identify *Mtb*-reactive CDR3αβ sequences and characterize their functions. (L) Frequencies of CD69-expressing T cells, measured by flow cytometry after stimulation with PBS as negative control or *Mtb* lysate. (M) Schematic representation of the migration and activation of *Mtb*-reactive T_EM_ cells within granulomas.

The spatial proximity of *LGALS9*^high^ macrophages and T_EM_ cells lead us to investigate whether infiltrating CD8⁺ T cells recognize *Mtb*-derived antigens locally. Among the CD8⁺ T-cell populations, CD8_PDCD1/CCL4 T_EM_ cells showed the strongest activation signatures, including enrichment of TCR signaling and chemokine-response pathways (**Figure 4E**). These cells also expressed effector genes such as *IFNG* and *TNF*, while simultaneously expressing exhaustion-associated markers (**Figure 4F**). TCR repertoire analysis revealed a significant increase of clonal expansion for these T_EM_ cells in TB samples (**Figures 4G, S8B and S8C**). Expanded clones preferentially expressed cytotoxic molecules (*GZMH* and NKG7), activation-associated genes (*HLA-DRB1*, *HLA-DPA1*, *HLA-DRB5*), and memory-associated genes (*THEMIS* and *LGALS1*) (**Figure S8D**), consistent with antigen-driven activation. To assess potential antigen specificity, we used TABR-BERT^70^ to predict TCR recognition of immunodominant *Mtb* antigens, including PE, PPE, and ESAT-6 proteins. A subset of T_EM_ cells had higher predicted antigen-recognition scores than other T-cell populations (**Figure 4H**), supporting the presence of expanded, potentially *Mtb*-reactive effector cells in granulomas.

Because granuloma development and maintenance can involve recruitment of T cells from distal tissue regions^71^, we next asked whether activated T_EM_ cells were shared between distal regions and lung lesions. TCR-based lineage tracing suggested that more than half of the T_EM_ cells in TBL were related to clones detected in TBD (**Figure 4I**). In contrast, inferred sharing was limited for most CD4⁺ T-cell, with T_H_1 cells as an exception, and for B-cell subsets (**Figure S8E**). Compared with non-migratory counterparts, migratory T_EM_ cells expressed higher levels of activation-associated genes (*HLA-DRB1*, *HLA-DRB5*, and *COTL1*) and the cytotoxic effector gene *GZMH* (**Figure 4J**). These findings are consistent with preferential recruitment of activated, antigen-experienced CD8^+^ T cells into granulomas. Spatial analysis showed that T-cell activation signatures were concentrated around granulomas, whereas exhaustion and senescence signatures were highest within SD6 (**Figures S8F and S8G)**. This pattern suggests that persistent antigen exposure may drive functional exhaustion after CD8^+^ T cell infiltration. Nevertheless, these cells may retain effector potential, including the capacity to produce multiple cytokines, while exhibiting reduced cytotoxicity, particularly during or shortly after recruitment. This functional state resembles the rheostat-like phenotype described for HBV-specific CD8^+^ T cells during chronic HBV infection^72^.

To functionally assess the predicted antigen reactivity, we generated 264 TCR constructs from migratory T_EM_ cells with the highest TABR-BERT prediction scores from individuals carrying HLA-A*02:01 or HLA-A*24:02 (**Figure 4K**). TCR-engineered T cells were co-cultured with HLA-matched K562 antigen-presenting cells pulsed with *Mtb* lysate. Compared with PBS controls, *Mtb* lysate stimulation increased the frequency of activated T cells (**Figure 4L**). These results provide functional evidence that the computationally prioritized migratory CD8^+^ T_EM_ repertoire is enriched for TCRs responsive to *Mtb* lysate-presenting cells.

Together, these findings identify a macrophage-T_EM_ niche associated with the local recruitment and activation of *Mtb*-reactive CD8⁺ T cells by *LGALS9*^high^ chemokine expressing macrophages and may contribute to the organization of adaptive immunity within granulomas (**Figure 4M**).

### *LGALS9*^high^ chemokine-expressing macrophages are required for adaptive immune cell recruitment into granuloma for bacterial control in zebrafish

We next ask whether *LGALS9*^high^ macrophages organize adaptive immune niches during granuloma development and whether these niches contribute to bacterial control. Given the antibacterial transcriptional program of *LGALS9*^high^ macrophages, we first examined whether this population was enriched in granulomas with effective bacterial containment *in vivo*. Previous work^24^ in macaques showed that, at 10 weeks after infection, granulomas with lower bacterial burdens were associated with enhanced antimicrobial activity. We therefore compared the transcriptional profiles of granulomas with low and high bacterial burdens at 10 weeks post infection using data from Gideon et al.^24^ Genes significantly upregulated in low-burden granulomas were defined as a low-bacterial-burden signature, representing transcriptional programs associated with bacterial containment (**Figure 5A**). Among all the myeloid populations in our scRNA-seq data, *LGALS9*^high^ macrophages (AM_LGALS9/CXCL10) and DC_LAMP3 showed the strongest enrichment for this signature (**Figure 5B**). The corresponding spatial domains, SD6 and SD10, were also strongly enriched (**Figure 5C and S9A**). These findings indicated a significant intrinsic correlation between the activated alveolar macrophages and granulomas with low bacterial burdens.

**Figure 5.**
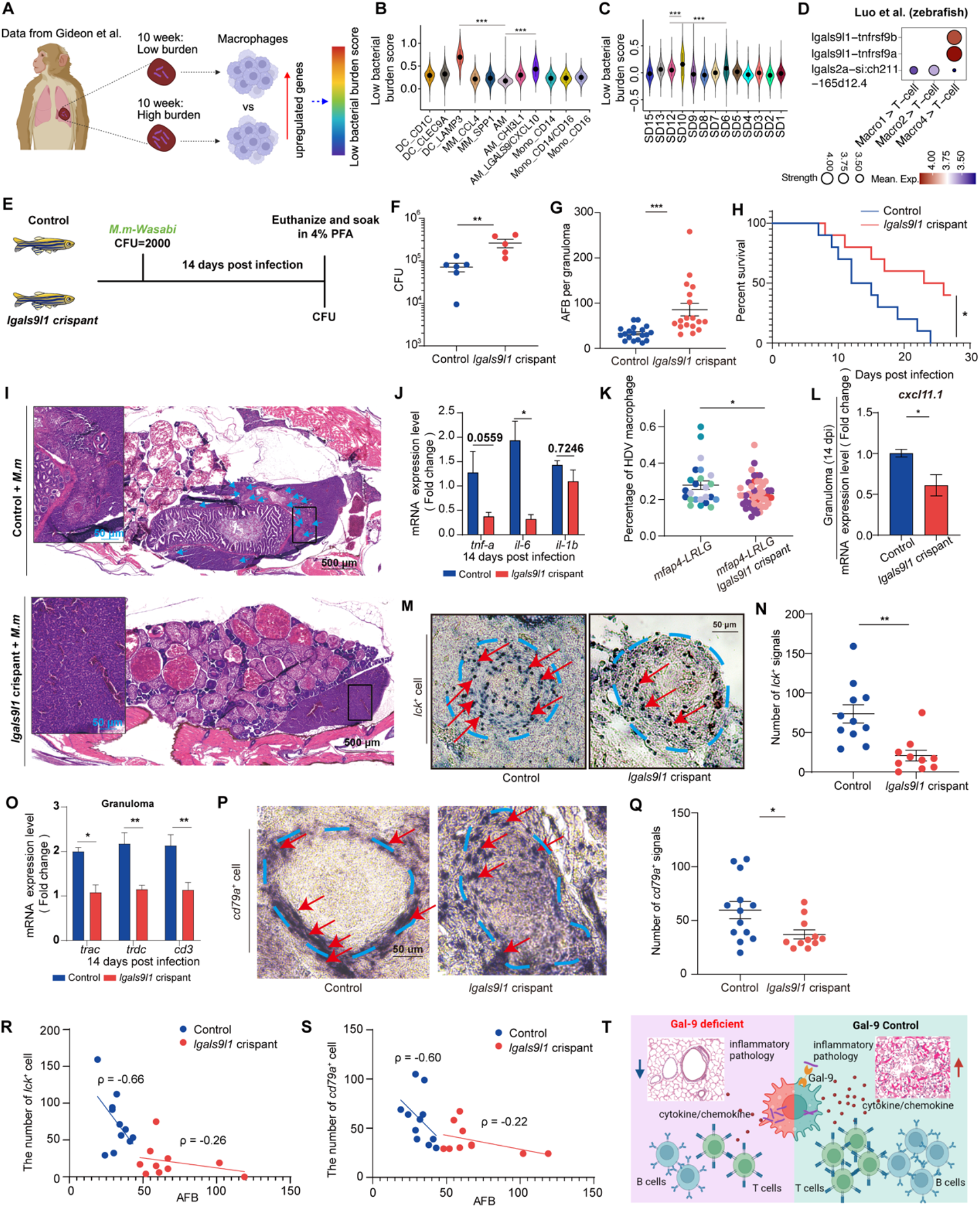
*LGALS9*^high^ chemokine-expressing macrophages correlated with bacterial containment and adaptive immune niche formation. (A-C) Definition of low bacterial burden scores (A) and their distribution across myeloid populations (B) and spatial domains (C) in macaque granulomas. Only data from the late stage (10 weeks) after infection were included. (Mann-Whitney U Test, \*\*\**P* < 0.001). (D) Macrophage-T cell communication inferred from zebrafish granuloma scRNA-seq data. (E) Schematic representation of the experimental design used to assess *lgals9l1* knockout in zebrafish infected with *Mycobacterium marinum*. (F) Bacterial load in zebrafish from the control (Ctrl) and *lgals9l1* crispant groups. Each point represents the bacterial load in an individual zebrafish from one biological replicate. (G) Acid-fast bacilli (AFB) count in granulomas from the Ctrl and *lgals9l1* crispant groups. Each point represents the AFB count in an individual granuloma. (H) Survival curves for zebrafish from the Ctrl and *lgals9l1* crispant groups. Each group contained 10 fish. (I) H&E staining of paraffin-embedded zebrafish sections from the Ctrl and *lgals9l1* crispant groups. Blue arrows indicate granulomas. (J) Expression of key inflammatory cytokines in the Ctrl and *lgals9l1* crispant groups. (K) Proportion of HDV macrophages in the Ctrl and *lgals9l1* crispant groups. Each point represents the statistical value for an individual granuloma. Different colors indicate three independent biological replicates, with data obtained from three zebrafish per experiment (each zebrafish having 2 to 3 granulomas). (L) The mRNA expression level of *cxcl11.1* in the Ctrl and *lgals9l1* crispant groups. (M) Representative images of *lck+* in situ hybridization signals in granulomas from the Ctrl and *lgals9l1* crispant groups. Blue dashed lines outline the granulomas, and red arrows indicate *lck+* in situ hybridization signals. (N) Quantification of *lck+* in situ hybridization signals in different granuloma types from the Ctrl and *lgals9l1* crispant groups. Each point represents the number of *lck+* signals in an individual granuloma. (O) Relative mRNA expression levels of the TCR-related genes *trac, trdc* and *cd3* in granulomas from the Ctrl and *lgals9l1* crispant groups. Each group consists of three zebrafish. (P) Representative images of *cd79a+* in situ hybridization signals in granulomas from the Ctrl and *lgals9l1* crispant groups. Blue dashed lines outline granulomas, and red arrows indicate *cd79a+* signals. (Q) Quantification of *cd79a+* in situ hybridization signals in different granuloma types from the Ctrl and *lgals9l1* crispant groups. Each point represents the number of *cd79a+* signals in an individual granuloma. (R-S) Correlations between AFB counts and the number of *lck*+ cells (R) or *cd79a+* cells (S) in individual granulomas from the Ctrl and *lgals9l1* crispant groups. (T) Schematic representation of the contrasting outcomes in the Ctrl and *lgals9l1* crispant groups. * *P* < 0.05, ** *P* < 0.01, *** *P* < 0.001.

To investigate the function of *LGALS9* and its associated signaling *in vivo*, we employed the zebrafish-*Mycobacterium marinum* infection model, which recapitulates several structural and immunological features of human tuberculous granulomas^73,74^. Consistent with our human findings, analysis of a previously generated zebrafish single-cell atlas^16^ using DanioTalk^75^ identified the *lgals9l1-tnfrsf9a/b* interaction as the strongest inferred macrophage-T-cell signaling axis (**Figure 5D**). This pathway represents a functional orthologue of the human LGALS9-TNFRSF9 pathway (**Table S6**). We generated *lgals9l1* deficient zebrafish using CRISPR-Cas9 (**Figure 5E**). Loss of *lgals9l1* did not significantly alter bacterial burden in infected embryos which do not have adaptive immunity (**Figure S9B**). In contrast, adult *lgals9l1* crispants had markedly increased bacterial loads at 14 days post-infection (**Figures 5F, 5G** and **S9C**), suggesting that *lgals9l1* contributes to bacterial control during mature immune responses. Despite their higher bacterial burdens, mutant fish survived significantly longer than controls (**Figure 5H**) and showed reduced inflammatory pathology (**Figures 5I** and **5J**). Histological analyses further revealed a reduced macrophage accumulation within granulomas in *lgals9l1*-deficient fish (**Figures 5K** and **S9D**). Although zebrafish lack a direct CXCL10 orthologue, it expresses *cxcl11*, another IFN-γ-inducible CXCR3 ligand^76^, which was significantly lower in granulomas from *lgals9l1*-deficient animals (**Figures 5L**), consistent with impaired chemokine-mediated immune recruitment.

In human samples, spatial analysis showed that *LGALS9*^high^ macrophages formed interaction niches with T_EM_ cells and are located near B-cell-enriched regions. Consistent with this organization, CellChat^77^ analysis also identified spatial interactions between SD6/SD10 and SD1 (**Figure S9E**). The different outcomes in embryonic and adult infection models, particularly the lack of a significant bacterial-burden effect in embryos, prompted us to investigate whether *lgals9l1* contributes to adaptive immune-cell recruitment. In situ hybridization showed markedly fewer *lck*⁺ T cells within granulomas of *lgals9l1* crispants (**Figures 5M** and **5N**), accompanied by significantly reduced *tcrα* and *cd3* transcript levels (**Figure 5O**). Recruitment of *cd79a⁺* B cells were also substantially reduced in mutant granulomas (**Figures 5P** and **5Q**). In control animals, bacterial burden negatively correlated with the abundance of intra-granulomatous T and B-cells (**Figures 5R** and **5S**); these correlations were markedly attenuated in *lgals9l1* crispants. Together, these results suggest that *lgals9l1* supports the recruitment and organization of adaptive immune cells within granulomas and may thereby contribute to bacterial containment.

Collectively, these findings support a model in which *LGALS9*^high^ chemokine-expressing alveolar macrophages establish localized immune niches that link innate bacterial sensing to the recruitment and organization of adaptive immune cells (**Figure 5T**). Through *LGALS9*-dependent signaling and chemokine networks, these macrophages may promote lymphocyte accumulation within granulomas, thereby supporting local antigen-specific responses while influencing the balance between bacterial control and inflammatory pathology. Thus, *LGALS9*^high^ macrophages may help organize granuloma-associated adaptive immunity in addition to function as antimicrobial effector cells.

### Tuberculosis severity is associated with spatial remodeling of TLSs

Given the potential role of these two adaptive immune niches in bacterial containment, we next investigated their spatial relationship with disease severity in human TB. We trained the STflow^78^ model using spatial transcriptomic datasets from six patients, using 330 genes selected from differentially expressed genes of SD1 and SD6 (**Figure 1A**). Following five-fold cross-validation, TLS-associated marker genes, including *MS4A1* and *CXCL13*, showed strong and consistent spatial correlations between predicted and measured expression (**Figure 6A**), supporting the model’s ability to reconstruct TLS-associated transcriptional programs. In contrast, prediction of T_EM_-associated programs performed poorly (**Table S7**). We therefore focused subsequent analyses on T_FH_-B cell niches and predicted their spatial gene-expression profiles from H&E-stained sections of 137 TB samples. Patients were stratified into mild and severe TB groups according to predefined criteria integrating radiological findings, surgical indications, and pathological diagnosis (**Methods**; **Table S8**). The cohort included 37 patients with mild TB and 44 patients with severe TB, all of whom recovered after surgery. Where available, multiple sections containing distinct granulomas from the same patient were included in the analyses; each section represented a single granuloma.

**Figure 6.**
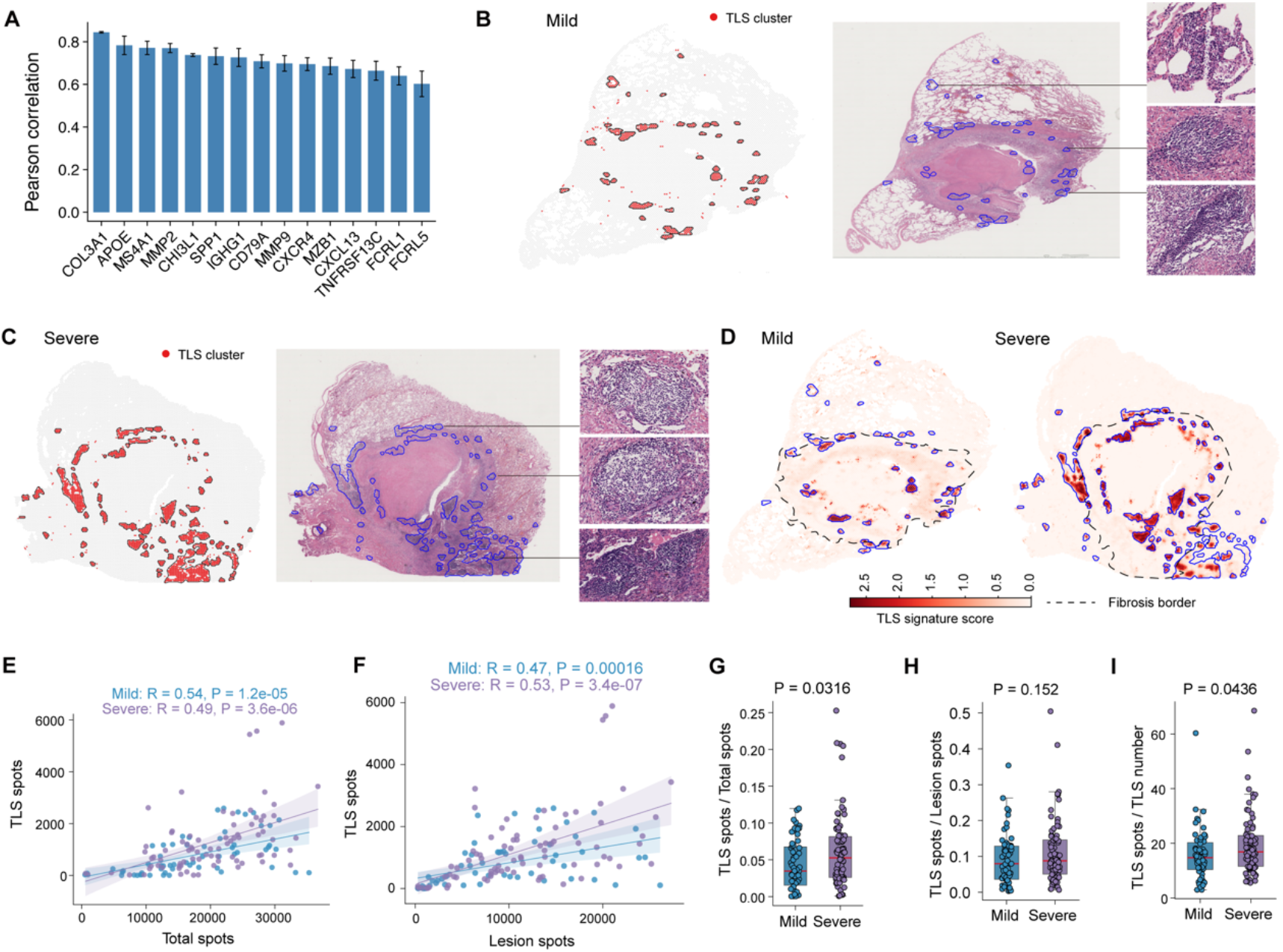
TLS abundances are associated with TB disease severity. (A) Pearson correlation between predicted and observed gene expression levels in spatial transcriptomic data. (B) Representative slide from a mild TB case, with identified TLSs highlighted on the H&E-stained section and enlarged views of representative TLS structures. (C) Representative slide from a severe TB case, with identified TLSs highlighted on the H&E-stained section and enlarged views of representative TLS structures. (D) Spatial distribution of TLS signature scores on representative mild and severe TB sections. (E) Spearman correlation between the total number of spots and the number of TLS spots. Each point represents one TB sample. (F) Spearman correlation between the number of lesion spots and the number of TLS spots. Each point represents one TB sample. (G) Ratio of TLS spots to total spots in mild and severe TB cases. Each point represents one TB sample (Two-tailed t-test). (H) Ratio of TLS spots to lesion spots in mild and severe TB cases. Each point represents one TB sample (Two-tailed t-test). (I) Ratio of TLS spots to the number of TLSs in mild and severe TB cases. Each point represents one TB sample (Two-tailed t-test).

Unsupervised clustering of the predicted transcriptional profiles identified cluster 6 carrying a characteristic TLS-associated signature (**Figures S10A** and **S10B**). We therefore defined spatially contiguous spots assigned to cluster 6 as individual TLSs, requiring a minimum of three spots per TLSs. The number of spots within each TLS was used as a measure of TLS size. The identified TLS niches were also confirmed by immumohistochemical staining in mild patients (**Figures 6B**) and severe patients (**Figures 6C**). To further validate these niches, we calculated a TLS signature score based on *MS4A1*, *CXCL13* and *FCRL3*. Spatial mapping of this score confirmed the TLS niches identified on H&E-stained sections in both mild and severe patients (**Figure 6D**). Consistent with the spatial transcriptome data, the predicted TLS niches were distributed along the fibrotic boundaries of granulomas as inferred using RCTD^79^ (**Figures 6D, S10C** and **S10D**).

TLS niches were more abundant in severe TB samples (**Figures 6B-D**). However, this difference could have been influenced by variation in granuloma size. To minimize this potential confounding factor, we identified lesion clusters by matching lesion-associated domains from the spatial transcriptomic data to virtual clusters using RCTD (**Figure S10E**). As expected, the lesion regions are located within the fibrotic boundaries of granulomas (**Figure S10F**). TLS abundance was positively correlated with both the total number of spots across the entire tissue section and the number of spots number within lesion regions (**Figures 6E** and **6F**). Nevertheless, after normalization to the total number of spots per section, TLS abundance remained significantly higher in severe TB than in mild TB (**Figure 6G**). A similar trend was observed after normalization to lesion size, although the difference did not reach statistical significance (**Figure 6H**).

The ratio of total TLS size to TLS number, representing mean TLS size, was also significantly higher in severe TB (**Figure 6I**), indicating larger TLS niches in severe disease. However, the TLS maturation score, calculated as the mean inferred expression of the established maturation markers *CR2* and *FCER2*, which encode CD21 and CD23, respectively^80^, did not differ significantly between mild and severe TB (**Figures S10G**-**S10I**). Together, these findings associate TB severity with changes in TLS abundance, without corresponding evidence of increased TLS maturation based on the inferred transcriptional profiles. Such remodeling may reflect persistent antigen exposure and sustained inflammatory recruitment; its relationship to protective immunity remains to be established.

## Discussion

Granulomas are dynamic immunological structures in which *Mtb* persistence and host defense are spatially intertwined^36^. Although traditionally viewed as sites of immune containment involving macrophages and lymphocytes^2^, granulomas are increasingly recognized as heterogeneous microenvironments in which distinct immune populations may exert divergent effects on bacterial control and tissue pathology. By integrating single-cell transcriptomic, spatial and immune-receptor data, we provided an unprecedented granulomas’ spatial-structure at single-cell resolution and identify two spatially distinct adaptive immune niches around the necrotic core. The outer T_FH_-B cell niche has active TLS function and contains sequential B cell states consistent with differentiation toward antibody-producing PCs, suggesting that these TLSs may support antigen-driven B-cell maturation beyond their previously described architectural role. The inner CD8⁺ T_EM_-macrophage niche correlated with the accumulation of antigen-experienced, potentially *Mtb*-reactive CD8^+^ T cells surrounding the necrotic core. *LGALS9*^high^ chemokine-expressing macrophages are spatially associated with these niches, while zebrafish *lgals9l1* perturbation implicates this pathway in adaptive immune-cell accumulation and bacterial control. Thus, rather than serving solely as sites of immune containment, granulomas may function as spatially organized adaptive immune hubs in which specialized niches coordinate humoral and cellular immunity during persistent *Mtb* infection.

TB granulomas have complex cellular composition and spatial organization, but the mechanisms underlying their formation and function remain poorly understood. Histologically, they comprise macrophage-rich cores surrounded by lymphocytes and fibroblasts^10,12^. Single-cell and spatial profiling, together with studies across diverse animal models, has advanced our understanding of TB granuloma biology; however, the direct understanding of granuloma organization and immune responses in human TB remains limited^14,18,23,24,81^. Recent human spatial studies have revealed layered architectures and specialized niches, but have been limited by small marker panels and have provided only partial insights into the complex immune microenvironment^3,31,34^. Here, we create a resource for characterizing granuloma spatial organization and local immunity at single cell resolution (https://tbomics.smart-nbc.org.cn/). Compared with previous studies, this resource combined transcriptomic and spatial data with paired TCR and BCR repertoire profiles and includes lesional, adjacent and distal lung tissues, as well as healthy lung as controls. We also profiled tonsil tissue as a reference for spatial TLS organization. These data enable characterization of adaptive immune populations and their associated molecular niches in tuberculoma-associated TLS. More importantly, the paired TCR and BCR repertoires in this study may especially help future studies identify antigen-specific receptors, antibodies, and relevant immunodominant B-cell antigens and T-cell epitopes through functional validation. Therefore, this comprehensive single-cell atlas of human granuloma provided a foundation for understanding granuloma adaptive immunity and particularly facilitating the future development of host-directed therapies and vaccines for TB (https://tbomics.smart-nbc.org.cn/)

Basing on this resource, our study also shows spatial evidence that *Mtb*-reactive CD8⁺ T_EM_ cells accumulate in the inner region of human granulomas, surrounding the necrotic core and in proximity to tissue-resident alveolar macrophages. These macrophages have been reported to be frequent targets of *Mtb* and to express infection-associated and immune-migratory programs^82^. TCR-engineered cells bearing selected receptors from migratory CD8^+^ T_EM_ cells responded to *Mtb* lysate, providing the functional evidence that the prioritized repertoire is enriched for *Mtb*-responsive TCRs. This assay does not, however, establish the specificity or cognate epitopes of every predicted receptor. We observed two spatially distinct transcriptional states among migratory CD8^+^ T_EM_ cells. At the periphery of granulomatous regions, these cells displayed stronger signatures of TCR activation, cytotoxicity and effector differentiation than their non-migratory counterparts. Within granulomatous lesions, exhaustion– and senescence-associated signatures were more predominant. This pattern is consistent with a dynamic state in which continuous replenishment and functional exhaustion coexist. Persistent expression of *IFNG*, *TNF* and cytotoxic-associated genes indicate that these cells may retain some effector potential despite chronic stimulation, although transcript expression alone does not establish cytokine polyfunctionality or cytotoxic capacity. Continuous recruitment from distal lung compartments replenish the local pool of antigen-experienced cells, but this model requires direct testing. This interpretation is consistent with recent work in chronic HBV infection showing that tissue-specific cAMP-PKA signaling can restrain virus-specific CD8⁺ T-cell activation and effector function through a rheostat mechanism distinct from canonical T-cell exhaustion^72^. It highlights how persistent antigen exposure and local tissue cues can shape CD8⁺ T-cell states beyond a simple functional-versus-exhausted dichotomy. However, whether a similar mechanism operates in TB granulomas remains unknown. Our finding may have therapeutic implications, as residual effector CD8^+^ T_EM_ cells within granulomas could potentially be reactivated by renewed antigen exposure such as therapeutic vaccine. In light of evidence that CD8^+^ T cells are required for early protection against *Mtb*^27^, future work should more comprehensively define the *Mtb* antigen specificity of these migratory CD8^+^ T_EM_ cells and identify dominant epitopes presented by different HLA molecules.

The outer niche reveals a distinct form of adaptive immune organization centered on T_FH_-B cell interactions within TLSs^34,43^. These structures may provide a local environment for B-cell selection and differentiation. Although previous studies have described GC-like features in granuloma-associated TLSs^32^, our data identify the associated B-cell subpopulations and suggest a potential trajectory of B-cell states. B_FCRL3, B_EGR1, and B_FCRL5 exhibit transcriptional features associated with GC-like activation, class switching, somatic hypermutation, and PC differentiation. These findings are consistent with ongoing antigen-driven B-cell maturation rather than simply passive accumulation. Along the inferred trajectory, IgM^+^ B_FCRL3 cells show dark-zone-like features and may represent an earlier state, whereas B_FCRL5 cells showed atypical memory B-cell features alongside programs associated with GC activity and PC differentiation^83^. The function of FCRL3^+^ B cells and their contribution to TLS formation and maturation in granulomas require further investigation.

Granuloma-associated TLSs shared the similar spatial features with tonsillar GCs, while they exhibit a thicker IgD⁺ B-cell mantle and greater accumulation of surrounding PCs. These differences may reflect sustained local antigenic stimulation^84–86^ and provide additional evidence of local antibody generation in lung. B cells may also organize T_FH_-like cells within TLSs and contribute to *Mtb* control^45^. Given the phenotypic diversity of TLS-associated memory B cells, functional studies are needed to determine which differentiation programs or B cell populations are protective, maladaptive, or context-dependent during human TB. Lung antibodies have been reported to react with *Mtb* and promote its phagocytosis^42^, while patients with severe TB have increased numbers of PCs^87^. These observations support a potential role for local humoral responses in *Mtb* containment. However, the positive association between TLS abundance and disease severity in our cohort indicates that increased lymphoid organization does not necessarily correspond to effective bacterial control. This contrasts with animal-model studies in which appropriately organized B-cell responses contribute to early bacterial containment^45,47,52^. Differences in disease stage may partly explain this discrepancy. The functional state and spatial organization of B cells may therefore be more informative than their abundance alone. Why patients with TB, particularly those with severe disease, failed to control *Mtb* despite abundant pulmonary PCs remains an important question.

We identified an underappreciated *LGALS9*^high^ macrophage population in human granulomas. This population shares features with previously described TB-susceptible tissue-resident alveolar macrophages^82^ but is distinct from monocyte-derived pro-inflammatory macrophages and *SPP1*^+^ macrophages^15^, which can interact with MMP1^+^CXCL5^+^ fibroblasts in the granuloma cuff and contribute to immunopathology^88^. Our findings extend the known heterogeneity of granuloma-associated macrophages. Galectin-9 (Gal-9), encoded by *LGALS9*, can recognize mycobacterial arabinogalactan, an essential component of the *Mtb* cell wall, through its carbohydrate-recognition domain 2^89^. In macaques with late-stage *Mtb* infection, *LGALS9*^high^ macrophages were associated with low-bacterial-burdens in granulomas. In zebrafish, this macrophage population was associated with *lgals9l1*⁺ macrophages^16^, and *lgals9l1* deficiency significantly increased bacterial burden in adult animals. Our findings further suggest *LGALS9*^high^ chemokine-expressing macrophages regulate the formation of adaptive immune niches and inflammatory response within granulomas.

These macrophages were spatially associated with T_EM_ cells through CXCL9/10/11-CXCR3 and LGALS9-associated pathways. Consistent with this observation, T-cell and B-cell accumulation within granulomas was substantially reduced in *lgals9l1*-deficient zebrafish. This impairment of adaptive-cell recruitment was accompanied by reduced inflammation and prolonged survival despite increased bacterial burden, which is consistent with the observation that higher *Mtb* loads are often associated with attenuated inflammation during the establishment of chronic infection^90,91^. Together, these findings support a model in which macrophage-derived chemotactic and costimulatory signals maintain a local adaptive immune response that contributes to bacterial containment while also participating in inflammatory pathology. However, because the zebrafish perturbation was systemic, the extent to which these effects are macrophage intrinsic remains to be determined.

Our study also highlights the utility and limitations of AI-based spatial gene-expression prediction from H&E-stained sections. Image-based prediction of spatial gene expression has become increasingly useful^92,93^, particularly in cancer research^93–95^. We trained a TB-specific model using spatial transcriptomic data with 55-μm spots. The model predicted TLS-associated programs with reasonable accuracy but performed less well for the macrophage-T_EM_ niche. The limited resolution of the training data, with approximately 1∼10 cells per spot, may limit the model’s ability to resolve individual T_EM_ cells and their interactions with macrophages. The small training cohort also limits generalizability. Although TLS size may vary with abundance and maturation, the current data could not resolve these processes in sufficient detail. Higher-resolution spatial profiling and larger TB-specific training datasets will be needed to identify spatial niches associated with disease progression, bacterial control, and protection.

In summary, our findings identify two adaptive immune niches that are interconnected components of a spatially organized innate-adaptive immune circuit. Antigen-experienced T cells and B-cell responses are integrated into granuloma architecture alongside local myeloid cells. H&E-based spatial transcriptomic reconstruction revealed increased TLS abundance in severe TB, without evidence of increased maturation based on the inferred transcriptional profiles. This remodeling may reflect persistent antigen exposure and the accumulation of functionally heterogeneous lymphoid niches, but its contribution to clinical outcomes remains unsolved. Integrating spatial organization with cellular function, antigen specificity and clinical outcome will be essential for distinguishing protective from dysfunctional immune structures and for guiding the development of host-directed therapies and vaccines against TB.

### Limitations of study

The ability to determine how these spatial niches relate to controlled or progressive TB was limited by cohort size, tissue availability, and sampling constraints. Larger, longitudinal cohorts with broader sampling of granulomas and distal lung tissue will be needed to establish whether the identified niches are associated with disease control, progression, treatment response, or specific clinical outcomes. Given the limited sample size, this study does not support reliable stratified comparisons among different lesion types, such as caseous, fibrotic, and calcified lesions. Experimental perturbation studies will be required to determine whether *LGALS9*^high^ macrophages directly regulate the formation and function of the T_EM_ and TLS-associated niches. Although our multi-omics analyses support antigen-driven responses within granulomas, the antigen specificity of T_FH_ and several TLS-associated B cell populations remain to be validated experimentally. Higher-resolution spatial technologies, complementary imaging approaches, and larger cohorts will be needed to determine how the immune cell subpopulations are organized and how they interact with the niches identified in this study.

## Data availability

- The sequencing raw data used in this paper are available in GSA (Genome Sequence Archive in the BIG Data Center, Beijing Institute of Genomics, Chinese Academy of Sciences) under accession numbers HRA003733. The processed raw count matrices are available in our TB atlas website (https://tbomics.smart-nbc.org.cn/). The code performing data analysis and figure drawing was available upon request.
- All software and analytical methods used in this study are publicly available.

## Acknowledgement

This study was supported by the National Natural Science foundation of China (82625106 to Z.Z., 82372263 to B.Y.), the National Key R&D Program of China (2023YFC2308300 to F.Q.), the R&D Program of Guangzhou Laboratory (SRPG22-006 to Z.Z.), the Shenzhen Science and Technology Program (ZDSYS20210623091810030 to Z.Z., KQTD20200909113758004 to Z.Z.) and the Shenzhen Clinical Research Center for Tuberculosis (20210617141509001 to Z.Z.).

## Author contributions

All authors had full access to the data in the study and accepted responsibility to submit it for publication. Conceptualization: Z.Z. and F.Q.; Methodology: Z.Z., F.Q., Y.C., Q.Y., B.Y., H.W., W.Z. and Y.Y.; Software: F.Q., Y.C., R.H., Y.Y.; Validation: Q.Y., H.W., W.Z., G.L.; Formal analysis: F.Q., Y.C., Q.Y., R.H.; Investigation: F.Q., Q.Y., H.W., and T.R.; Resources: K.Q., Z.C., M.L., T.R., J.Z.; Data curation: K.Q. T.R., Z.C., M.L., G.L., J.Z.; Funding acquisition: Z.Z., F.Q., B.Y; Project administration: Z.Z. and F.Q.; Supervision: Z.Z., F.Q. and B.Y.; Visualization: F.Q., Y.C., Q.Y., D.B., LM.N. and H.W.; Writing – Original Draft: Z.Z. and F.Q.; Writing – Review & Editing: all authors.

## Methods

Key resources.

| **REAGENT or RESOURCE** | **SOURCE** | **Catalog No. or Clone No.** |
|---|---|---|
| **Antibody** |  |  |
| CD3 | Biolynx | BP6027 |
| CD4 | abcam | ab252199, EPR6855 |
| CD8 | abcam | ab17147, C8/144B |
| CD20 | CST | 74332T, L26 |
| CD68 | Biolynx | BP6036 |
| CXCL13 | ZSGB-BIO | ZG-0601 |
| CXCR5 | CST | 72172, D6L3C |
| CXCL10 | Proteintech | 10937-1-AP |
| TIGIT | CST | 99567, E5Y1W |
| BCL-6 | abcam | ab172610, EPR11410-43 |
| CD10 | abcam | ab227640, SP67 |
| IgG | ZSGB-BIO | ZA-0448 |
| IgD | ZSGB-BIO | BP6134 |
| **Biological samples** |  |  |
| TB samples | Shenzhen Third People’s Hospital | N/A |
| **Critical commercial assays** |  |  |
| Multi-tissue Dissociation Kit | RWD | DSC-800 |
| Chromium Next GEM Single Cell 5' Reagent Kits v2 | 10x Genomics | Cat# 1000263 |
| Chromium Single Cell Human TCR Amplification Kit | 10x Genomics | Cat# 1000252 |
| Chromium Single Cell Human BCR Amplification Kit | 10x Genomics | Cat# 1000253 |
| Chromium Next GEM SingleCell Fixed RNA Sample Preparation Kit | 10x Genomics | Cat# 1000414 |
| Chromium Next GEM Single Cell 3’ Reagent Kits v3.1 | 10x Genomics | Cat# 1000121 |
| Visium CytAssist Spatial Gene Expression for FFPE, Human Transcriptome | 10x Genomics | Cat# 1000522 |
| **Deposited data** |  |  |
| Human scRNA-seq, snRNA-seq, and spatial transcriptomics datasets | This paper | The National Genomics Data Center, China National Center for Bioinformation (CNCB-NGDC): HRA003733 https://tbomics.smart-nbc.org.cn/ |
| Zebrafish scRNA-seq data | Luo et al.<sup>16</sup> | The National Genomics Data Center, China National Center for Bioinformation (CNCB-NGDC): PRJCA027270 |
| Macaque scRNA-seq data | Gideon et al.<sup>24</sup> | Gene Expression Omnibus: GSE200151 |
| **Experimental models** |  |  |
| Zebrafish | Shanghai Public Health Clinical Center, Fudan University | N/A |
| **Software and algorithms** |  |  |
| CellRanger 6.1.1 | 10x Genomics | https://support.10xgenomics.com/single-cell-gene-expression/software/overview/welcome |
| SpaceRanger 3.0.0 | 10x Genomics | https://www.10xgenomics.com/support/software/space-ranger/downloads |
| Seurat 4.0.4 | Hao et al.<sup>96</sup> | https://satijalab.org/seurat/ |
| STARTRAC 0.1.0 | Zhang et al.<sup>97</sup> | https://github.com/Japrin/STARTRAC |
| COMMOT 0.0.3 | Cang et al.<sup>98</sup> | https://github.com/zcang/COMMOT |
| STRIDE 0.0.2 | Sun et al.<sup>50</sup> | https://github.com/DongqingSun96/STRIDE |
| MNMST 1.1 | Wang et al.<sup>49</sup> | https://github.com/xkmaxidian/MNMST |
| ROGUE | Liu et al.<sup>99</sup> | https://github.com/PaulingLiu/ROGUE |
| SCIGA | Ye et al.<sup>100</sup> | https://github.com/sciensic/SCIGA |
| Vegan 2.5.6 | Oksanen et al.<sup>101</sup> | https://github. com/vegandevs/vegan |
| STARTRAC 0.1.0 | Zhang et al.<sup>97</sup> | https://github.com/Japrin/STARTRAC |
| Monocle3 1.3.1 | Cao et al.<sup>66</sup> | https://cole-trapnell-lab.github.io/monocle3/docs/introduction/ |
| ggplot2 3.5.1 | Ginestet et al.<sup>102</sup> | https://ggplot2.tidyverse.org/ |
| Change-O 1.3.3 | Gupta et al.<sup>103</sup> | https://changeo.readthedocs.io/en/stable/ |
| SHazaM 1.0.2 | Gupta et al.<sup>103</sup> | https://shazam.readthedocs.io/ |
| msa 1.34.0 | Bodenhofer et al.<sup>104</sup> | https://github.com/UBod/msa |
| Alakazam 1.0.2 | Gupta et al.<sup>103</sup> | https://alakazam.readthedocs.io/ |
| CellChat 2.1.2 | Jin et al.<sup>77</sup> | https://github.com/jinworks/CellChat?tab=readme-ov-file |
| Scanpy 1.10.4 | Wolf et al.<sup>105</sup> | https://scanpy.readthedocs.io/en/stable/ |
| Python 3.10.9 | Python Software Foundation | https://www.python.org/ |
| R 4.3.3 | R Core Team | https://www.r-project.org/ |
| STFlow | Huang et al.<sup>78</sup> | https://github.com/Graph-and-Geometric-Learning/STFlow |
| clusterProfiler 4.0 | Wu et al.<sup>106</sup> | https://github.com/YuLab-SMU/clusterProfiler |
| NICHES | Raredon et al.<sup>107</sup> | https://github.com/msraredon/NICHES |

## Patient samples

This study was conducted in accordance with the principles of the Declaration of Helsinki. Ethical approval was obtained from the Research Ethics Committee of the Shenzhen Third People’s Hospital (China). Written informed consent was obtained from all participants. All experiments and sampling were performed in accordance with the ethical and biosafety protocols approved by our BSL-3 laboratory. All patients recruited in this study were diagnosed with active TB based on clinical symptoms, chest radiography, microscopy for acid-fast bacilli (AFB), sputum *Mtb* culture, and response to anti-TB chemotherapy. Granuloma biopsy samples were collected during surgery. Six lung tissues from non-diseased controls were obtained from deceased organ donors whose lungs were not used for lung transplantation.

A total of 9 active lesion tissues, 6 tissues from the lesion boundary or adjacent areas, and 9 tissues located distal to the lesions were collected and subjected to single-cell RNA (scRNA-seq) analysis. Additionally, 6 TB lesion tissues and 2 relative normal tissues were collected, which were utilized for spatial transcriptomics analysis and single-nucleus RNA (snRNA-seq) sequencing analysis. Furthermore, three formalin-fixed paraffin-embedded (FFPE) tissue blocks of TB lesion tissues were used for single and multiplex immunohistochemistry (mIHC) staining.

## Zebrafish studies

Zebrafish were maintained according to ethical guidelines (GB/T 35823-2018), with approval for experiments from the Animal Care and Use Committee of Shanghai Public Health and Clinical Center, Fudan University (2019-A016-01, 2022-A002-01, 2024-A20-01). All zebrafish strains had AB wild-type backgrounds. All zebrafish were reared under standard conditions^16^.

## 10x genomics scRNA-seq

Fresh tissues were subjected to digestion using the DSC-800 according to the manufacturer’s instructions. After 10 min of digestion at 37°C, the immune cells were purified using Percoll gradient centrifugation. The single-cell suspensions were washed twice with PBS and then resuspended with cell suspension solution at appropriate concentrations for subsequent scRNA-seq. The scRNA-seq libraries were prepared with the Chromium Next GEM Single Cell 5’ Reagent Kits v2 (10x Genomics) following the manufacturer’s instructions. Briefly, a total of 10 µL of single-cell suspension (containing ∼20,000 cells), 40 µL of barcoded gel beads, and partitioning oil were loaded onto Chromium Chip A/B to generate single-cell gel bead-in-emulsion. Polyadenylated transcripts were reverse-transcribed inside each gel bead-in-emulsion. Full-length cDNA along with cell barcode identifiers were PCR-amplified, and sequencing libraries were prepared and normalized. The constructed library was sequenced on a NoveSeq 6000 platform (Illumina, San Diego, CA).

## 10x genomics snRNA-seq

Nuclei isolation from FFPE tissue sections was performed according to the Chromium Fixed RNA Profiling Demonstrated Protocol (CG000632; 10x Genomics). Briefly, two 25-μm-thick sections were cut from each paraffin block and deparaffinized by sequential washes in xylene and graded ethanol solutions. Following a wash in ice-cold PBS, tissues were lysed in Lysis Buffer (Sigma-Aldrich) and mechanically homogenized using a pestle. The resulting pellet (containing released nuclei and undigested tissue) was resuspended, and isolated nuclei were filtered through a 20-μm cell strainer. Nuclei concentration and viability were quantified using a LUNA-FL™ Automated Cell Counter (Logos Biosystems, USA) after acridine orange/propidium iodide (AO/PI) staining; a subset was counterstained with DAPI for visualization. Qualified single nuclei were subsequently processed for Flex scRNA-seq library preparation.

Library synthesis was performed according to the Chromium Fixed RNA Multiplexed Samples protocol (CG000527; 10x Genomics) using the Chromium X platform. Following probe hybridization, uniquely barcoded samples were quantified and pooled at equimolar nuclei concentrations using the multiplexing strategy outlined in the Chromium Fixed RNA Profiling Multiplexed Samples Pooling Workbook (Rev B, CG000565). The pooled nuclei suspension was washed, and 20,000 nuclei per sample were targeted for partitioning into Gel Bead-in-Emulsions (GEMs) using a Chromium Chip Q. Subsequent steps included GEM barcoding, GEM recovery, cDNA pre-amplification, and library construction. Sequencing libraries were generated with the Chromium Next GEM Single Cell 3ʹ Reagent Kits v3.1 (10x Genomics, CG000315) following the manufacturer’s instructions. Final libraries were sequenced on an Illumina NoveSeq 6000 platform using paired-end dual-indexing (2 × 150 bp), achieving a median depth of approximately 300 million reads per sample.

## 10x genomics spatial transcriptomics

Consecutive sections were obtained from each human tissue FFPE block. These sections were stained to visualize tissue morphology and identify relevant regions of interest, and brightfield images were captured to document tissue orientation. Sections were then incubated with Visium Human Transcriptome Probes v2. These probes hybridize with specific mRNA transcripts and are linked to unique spatial barcodes corresponding to defined capture areas on the Visium slide. Each capture area measures 11×11mm and contains approximately 14,000 barcoded spots (55-micron diameter; 100-micron center-to-center distance). Following hybridization, probes are released from the tissue and bound to their complementary spatial barcodes on the slide. Captured mRNA were reverse transcribed into cDNA, and sequencing libraries were generated from this cDNA. Spatial barcodes and unique molecular identifiers (UMIs) were incorporated during library preparation to track transcript origin and individuality. Libraries were sequenced using a dual-index configuration on a NovaSeq 6000 platform (Illumina, San Diego, CA). The sequencing parameters were: 28 bp for Read 1 (containing a 16 bp Visium spatial barcode and 12 bp UMI), 50 bp for Read 2 (probe insert), 10 bp for the i7 sample index, and 10 bp for the i5 sample index. Paired-end sequencing was employed to capture both transcript sequences and spatial barcodes.

## H&E and immunohistochemistry staining

Segments of lung tissue from human TB patients who received surgery were collected and fixed in 10% buffered formalin (Sigma-Aldrich) and embedded in paraffin. Serial 5-mm sections of paraffin-embedded tissue were used for staining. Histologic sections were stained with hematoxylin and eosin (H&E) for pathological evaluation. Images of the whole microscope slide were captured using a NanoZoomer digital pathology system (Hamamatsu Photonics). For immunohistochemistry, tissues were fixed with acetone-chloroform for 3 minutes and then incubated for 2 hours with antibodies. Primary antibodies were detected using a biotinylated secondary antibody system following the manufacturer’s instructions. Images were captured using a 3DHISTECH digital pathology system (Pannoramic MIDI).

## Construction of gene knockout zebrafish

Guide RNAs (gRNAs) targeting *lgals9l1* (gRNA-T1: GGTATCTGTCCTGAGCGACA and gRNA-T2: GGAACCGCCTTACTATAACC) were designed using CRISPRscan (https://www.crisprscan.org/?page=gene). Scramble gRNA-T1 (CAGGCAAAGAATCCCTGCCG) and scramble sgRNA-T2 (TACAGTGGACCTCGGTGTCG) were designed as previously described (35545295). gRNAs and Cas9 protein (Novoprotein, E365-01A) were microinjected into zebrafish embryos at single cell stage. Genomic DNA was extracted at 24 hours post injection, and PCR was used to amplify the CRISPR site and the mutations were confirmed with Sanger sequencing. Details of gRNA sequences and primer sequences are:

gRNA sequence 1 forward primer: ATCTGAACAGGACCTGGGG

gRNA sequence 1 reverse primer: ACTCAATGGGGCTAATGCTG

gRNA sequence 2 forward primer: CCTGCTGTTATTTCAACTAAAC

gRNA sequence 2 reverse primer: GGGTTGACAACTCCATTG

## Bacterial strains

Single cell suspensions of *Mycobacterium marinum* M strain expressing wasabi fluorescent proteins were obtained as previously described^16^.

## Adult Zebrafish infection and bacterial load calculation

Adult zebrafish were intraperitoneally injected with approximately 2000 CFU of *Mycobacterium marinum*. Bacterial burden was determined by colony counting method. Briefly, infected zebrafish were euthanized with tricaine, followed by treatment with 75% ethanol and PBS for 3 minutes each. Zebrafish were then homogenized in a tissue homogenizer (Shanghai Jingxin Experimental Technology). Homogenates were treated with 4% NaOH for 10 minutes, gradient diluted, and plated onto 7H10 agar plates (containing hygromycin/amphotericin B/ polymyxin B/carbenicillin). Plates were incubated at 32°C for approximately 10 days, after which *Mycobacterium marinum* colonies was counted.

## Infection of Zebrafish Larvae

As previously decribed, at 3 days post fertilization, zebrafish embryos were infected via microinjection of ∼250 CFU into the duct of Cuvier, and bacterial burden was determined either by quantified bacterial fluorescence intensity or colony counting method^108^.

## Survival Curve

The death of zebrafish was recorded until all zebrafish succumbed, as previously described^16^.

## Paraffin Section In Situ Hybridization

Paraffin-embedded zebrafish sections were hybridized overnight at 65 °C with DIG-labeled *lck* antisense probes in hybridization buffer. Following hybridization, sections were incubated sequentially in 4× saline-sodium citrate (SSC) and 50% formamide in 2× SSC at 65 °C for 30 minutes, then treated with TNE buffer (Tris-HCl/NaCl/EDTA) at 37 °C for 10 minutes. Post-hybridization washes were performed with 2 × SSC and 0.2 × SSC at 65 °C for 15 minutes each. Sections were subsequently incubated overnight at 4 °C with anti-DIG-alkaline phosphatase antibody (Roche, 11093274910). Signal detection was carried out using NBT/BCIP (Roche, 11697471001) for at least 30 minutes, and the reaction was terminated once hybridization signals became clearly visible^109^.

## H&E staining of zebrafish tissue

Infected adult zebrafish were killed with tricaine, with their scales and fins removed, and immersed in 4% paraformaldehyde for at least 72 hours. Paraffin embedding, serial sectioning (4 to 5 µm), and HE staining were performed as previously described^110^

## Preparation of frozen section and antibody staining

The infected zebrafish were sacrificed with tricaine at 12 days post infection, and after scales and fins were removed, they were immersed in 1% paraformaldehyde and fixed at room temperature for 2 h. The samples were washed with PBS for 5 min, embedded with OCT, and then stored at –80°C. Serial sections (15-20 µm) were obtained for antibody staining. After the frozen sections were rinsed with PBS and pre-cooled with acetone, they were incubated with the blocking solution for 1-2 h. The sections were then incubated with the primary antibody (Takara 632496, 1:500 for DsRed) overnight at 4°C, and then incubated with the secondary antibody (Yeasen 34212ES60, 1:500 for Alexa Fluor 594) at room temperature for 2 hours. The sections were then sealed with an anti-fluorescence quencher^14,16^.

## Calculation of macrophage density in granuloma

To create a spatial distribution map of immune cells, we used Fiji software(v2.0) to convert the antibody staining signals on frozen sections into coordinate information (x and y)^111^. The midpoint of all coordinates was calculated (average x and y values) and normalized by dividing them at the midpoint. R software (v 4.3.1) was used to generate plots with standardized coordinates on a shared coordinate system and colors were added to the points based on their density, as previously described^110^. We defined density values greater than 90% as “high-density values (HDV)” and counted their proportions in each granuloma.

## RNA Extraction and Real-Time PCR

Whole zebrafish kidneys and granulomas from adults were homogenized through a 27G needle. cDNA was synthesized from 1 ug RNA using HiScript lll RT SuperMix for qPCR (Vazyme). Real-time PCR was performed with Hieff UNlCON Universal Blue qPCR SYBRGreen Master Mix (Yeasen) on a CFX96 Real-time system instrument (Bio-Rad).

## Gene Synthesis, Assembly, Cloning, and Sequence Verification

Synthetic oligonucleotides were chemically synthesized using the solid-phase phosphoramidite method on an automated DNA synthesizer. Oligonucleotides 60–120 nucleotides in length were designed to correspond to both the sense and antisense strands and to contain 15-25 bp overlaps to facilitate subsequent assembly. Following synthesis, all oligonucleotides were purified by denaturing polyacrylamide gel electrophoresis (PAGE) or high-performance liquid chromatography (HPLC) to remove truncated products and improve assembly efficiency. Full-length gene fragments were assembled from overlapping oligonucleotides by polymerase cycling assembly (PCA). Equimolar amounts of the oligonucleotides were combined in a single reaction, where they served as both primers and templates. Assembly reactions were performed using a high-fidelity DNA polymerase, such as Q5 High-Fidelity DNA Polymerase, Phusion High-Fidelity DNA Polymerase, Pfu DNA Polymerase, or KOD DNA Polymerase, through repeated cycles of denaturation, annealing, and extension. During thermal cycling, complementary overlapping regions annealed, enabling DNA polymerase to extend the oligonucleotides and progressively generate full-length double-stranded DNA. An aliquot of the initial assembly reaction was then used as the template for a second PCR with gene-specific outer primers to selectively amplify the complete gene fragment. The assembled DNA fragments were subsequently cloned into the target plasmid using Gibson Assembly. Briefly, the fragments and linearized vector were designed with homologous terminal sequences to facilitate assembly. Gibson Assembly reactions were performed using a commercially available master mix containing T5 exonuclease, a high-fidelity DNA polymerase, and DNA ligase, and were incubated at 50°C for 15–60 min according to the manufacturer’s instructions. During the reaction, T5 exonuclease resected the 5′ ends of the DNA fragments to generate complementary single-stranded overhangs, allowing homologous regions to anneal. DNA polymerase then filled the remaining gaps, and DNA ligase sealed the nicks to form intact recombinant plasmids. The recombinant plasmids were introduced into chemically competent Escherichia coli cells by heat shock. Transformed cells were plated on LB agar supplemented with the appropriate antibiotic-ampicillin, kanamycin, or chloramphenicol-according to the plasmid selection marker. Individual colonies were screened by colony PCR using vector-specific or gene-specific primers flanking the cloning site to confirm the presence and approximate size of the inserted fragment. Colonies testing positive by colony PCR were cultured overnight in selective LB medium, and plasmid DNA was purified using a commercial plasmid extraction kit according to the manufacturer’s instructions. Recombinant plasmids were initially validated by bidirectional Sanger sequencing to confirm the integrity of the entire insert. Sequencing reads were aligned with the designed reference sequence using standard sequence-alignment software, and only clones exhibiting 100% sequence identity were selected for downstream applications. For large-scale gene-library construction or the synthesis of long DNA fragments, pooled constructs were first characterized by next-generation sequencing (NGS), followed by isolation and sequence confirmation of individual positive clones by Sanger sequencing.

## Acid-fast staining (AFB)

Acid-fast staining was performed using the Kinyoun cold-staining method. Fixed smears were completely covered with carbolfuchsin solution (BASO, Zhuhai, China; Cat. No. BA4091A) and incubated for 5 min at room temperature without heating. The slides were gently rinsed with water and decolorized with acid alcohol (BASO; Cat. No. BA4091B) for 2 min, followed by another rinse with water. The smears were subsequently counterstained with methylene blue (BASO; Cat. No. BA4091C) for 30 s, rinsed with water, and air-dried. The stained smears were examined by bright-field microscopy using a 100× oil-immersion objective. Acid-fast bacilli were identified as red-stained rods against a blue background.

## H&E staining cohort and clinical classification

The H&E staining cohort consisted of patients with pathologically confirmed pulmonary tuberculosis who underwent pulmonary resection at Shenzhen third people’s hospital. Surgical intervention was performed for heterogeneous clinical indications, including therapeutic treatment of advanced pulmonary tuberculosis and diagnostic resection of indeterminate pulmonary lesions suspicious for lung malignancy. Therefore, tuberculosis severity was determined using integrated clinical criteria rather than surgical indication alone. Severe TB was defined by the presence of one or more of the following characteristics: (i) cavitary pulmonary tuberculosis, particularly multiple or enlarging cavities; (ii) destroyed lung or extensive pulmonary parenchymal destruction; (iii) radiological evidence of progressive pulmonary disease during follow-up; or (iv) major tuberculosis-related complications, including hemoptysis, pleural effusion, pneumothorax, or bronchiectasis associated with active tuberculosis. Mild TB was defined as localized pulmonary tuberculosis without cavitary disease, destroyed lung, or evidence of progressive active infection. Patients undergoing surgery primarily for diagnostic purposes, such as exclusion of lung malignancy, were also classified as mild TB when comprehensive evaluation of radiological evolution, microbiological findings, pathological diagnosis, and surgical indications showed no evidence of advanced pulmonary tuberculosis. To ensure cohort-wide consistency, all cases were re-evaluated using identical predefined criteria before downstream analyses. Because this cohort was derived exclusively from surgically resected pulmonary tissues, severity classification reflected the overall clinical status of pulmonary tuberculosis rather than the indication for surgery itself.

## Experimental validation of TCR candidates

### Construction of TCR-expressing J76.8 cells

Full-length TCRα-P2A-TCRβ chains of candidate TCRs were synthesized and cloned into the pHR_SFFV_IRES-EGFP lentiviral expression vector (GenScript). Lentivirus was produced by co-transfecting HEK293T cells in 6-well plates with the pHR vector containing the TCR insert, psPAX2 and pMD2.G. Viral supernatant was collected 72 h post-transfection and filtered through a 0.45 μM membrane (Millipore). J76.8 cells (5 × 10^5^) were seeded into 24-well plates and infected with 1.5 ml of viral supernatant at 700 × g for 1.5 h at 32 °C. After infection, 0.5 ml complete RPMI-1640 medium was added and cells were cultured at 37 °C for 3 days. TCR expression was assessed by flow cytometry after staining with PE anti-human CD3 (Biolegend, 344806), BUV805 anti-human CD8 (BD, 612889), APC anti-human TCRαβ (BD, 563826), and a viability dye (Thermo, 65-0866-18) for 30 min at 4 °C. The stained cells were washed twice with PBS and analyzed on a FACSymphony A3 (BD).

### Construction of K562-A*02:01 and K562-A*24:02 cells

A full-length HLA-A*02:01-or HLA-A*24:02-P2A-B2M construct was synthesized and cloned into pHR_SFFV_IRES-EGFP (GenScript). Lentivirus production and filtration were performed as above. K562 cells (5 × 10^5^) were seeded into 24-well plates and infected with 1.5 ml viral supernatant at 700 × g for 1.5 h at 32 °C. After infection, 0.5 ml of complete RPMI-1640 medium was added and cells were cultured at 37 °C for 3 days. HLA-A*02:01– or HLA-A*24:02 expression was quantified by flow cytometry after staining with APC anti-human HLA-ABC (Biolegend, 311410) and a viability dye (Thermo, 65-0866-18) for 30 min at 4 °C. The stained cells were washed twice with PBS and analyzed on a FACSymphony A3 (BD).

### Generation of TCR-T cells

Full-length TCRα-P2A-TCRβ fragments of candidate TCRs were excised from pHR_SFFV_IRES-EGFP with EcoRI and BamHI and subcloned into the SFG.MCS-3-mCherry retroviral vector. Retrovirus was produced by transfecting HEK 293T cells with the SFG vector and packaging plasmids (PegPam and RDF). Viral supernatant was collected 48-72 h later and filtered through a 0.45-μM membrane (Millipore). Human splenocytes were activated with T Cell TransACT (Miltenyi, 130-111-160) and cultured in GT-T551 medium (Takara, WK593S) supplemented with 5% heat-inactivated human AB serum (Gemini, 100-512) and 200 U/ml IL-2 (PeproTech, 200-02) for 4 days. For transduction, 1.5 ml viral supernatant per well was added to RetroNectin-coated (Takara, T100A) 24-well plates and centrifuged at 2000 × g for 1.5 h at 4°C to allow virus adsorption. After removal of viral supernatant, activated splenocytes (2 × 10^5^) per well were added and cultured in GT-T551 medium with 5% heat-inactivated human AB serum and 200 U/ml IL-2. Half of the medium was replaced every 3 days, and cells were expanded for 9 days before use in ICS assays.

### Activation of TCR-transduced J76.8 cells

TCR-transduced J76.8 cells (1 ξ 10^5^) were co-cultured with K562-A*02:01 or K562-A*24:02 (1 ξ 10^5^) in the presence of Mtb lysate in 96-well U-bottom plates. PBS treatment served as a negative control. NFAT-luciferase activity was measured after 8 h using the Bio-Lite Luciferase Assay System (Vazyme, DD1201-03). Cells were pelleted and resuspended in 50 μl PBS, followed by mixing with 50 μl substrate solution buffer for 5 min at room temperature before luminescence measurement was conducted on a Varioskan LUX plate reader (Thermo). In another assay, after 18h co-culture of TCR-expressing J76.8 cells and K562-A*02:01 or K562-A*24:02, CD69 expression was evaluated by staining with BV785 anti-human CD69 (Biolegend, 310932), and viability dye (Thermo, 65-0866-18) for 30 min at 4 °C, followed by PBS washes and analysis on a FACSymphony A3 (BD).

### Spatial Gene Expression Prediction from H&E Images

#### Image preprocessing

For each SVS whole-slide image, we used OpenSlide to access the image at pyramid level 0, thereby retaining the highest native spatial resolution available in the slide. The physical pixel size was determined primarily from the slide metadata. Based on the physical scale, we generated a staggered hexagonal grid across the entire slide and treated the grid points as pseudo-spots. By default, adjacent spot centers were separated by 100 μm, and each spot was represented by a square image region corresponding to a diameter of approximately 55 μm. To identify tissue-containing regions, we generated a thumbnail with a width of approximately 2048 pixels and constructed a tissue mask by combining grayscale intensity and color saturation information. Dark tissue regions were identified using Otsu thresholding, while visibly stained regions were retained based on their saturation values. The resulting mask was further refined using morphological closing and opening operations. Candidate spots were retained only when at least 20% of the corresponding image region was classified as tissue. For each retained pseudo-spot, a square image patch centered at the corresponding grid point was extracted from the highest-resolution image level. Regions extending beyond the slide boundary were filled with a white background. The extracted patches were resized to 224 × 224 pixels and normalized using the input statistics of the Hoptimus foundation model. The normalized patches were then processed by a vision Transformer, from which we extracted both the class-token representation, and the mean representation of the image patch tokens as image features for downstream spatial gene-expression prediction.

#### Training datasets

The six spatial transcriptomics (ST) datasets from TBL comprising paired high-resolution H&E-stained histological images and spot-level gene expression profiles were used to train models. Image preprocessing was conducted to align the histological features with the spatial transcriptomics coordinate space. For each spatial spot, image patches centered at the spot coordinates were cropped from the H&E images at 224×224 pixels. For the transcriptomic data, raw gene counts were preprocessed using Scanpy. The count matrices were normalized via log1p transformation to obtain the target ground-truth expression levels for model training.

#### Model Architecture and Training via STFlow

To predict spatial gene expression directly from histological morphology, we implemented the STFlow framework^78^. STFlow utilizes a flow-matching generative model that accounts for cell-cell interactions by modeling the joint distribution of gene expression across an entire slide. High-dimensional visual features are first extracted from cropped H&E image patches using a pathology foundation model and, together with spatial coordinates and iteratively refined gene-expression estimates, are used to condition the generation of spatial gene-expression profiles.

#### Cross-Validation and Evaluation

To rigorously evaluate the generalization performance of STFlow and prevent data leakage, a 6-fold cross-validation scheme was employed across the 6 ST datasets, where 5 samples were used for model training and hyperparameter tuning, while the remaining one sample served as the independent validation set in each iteration. The predictive performance was quantitatively assessed using the gene-wise Pearson Correlation Coefficient between the predicted gene expression profiles and the corresponding ground-truth spatial transcriptomics data. All models were trained using the AdamW optimizer with a learning rate of 0.0003 for 200 epochs.

### Cell type decomposition analysis

We used RCTD^79^ to map SD1-associated and fibrosis-associated transcriptional signatures onto predicted spatial gene-expression clusters using spatial transcriptomic and snRNA-seq data. The two analyses were performed independently. For the SD1-associated analysis, SD1-specific signature genes were used to define the reference expression profiles, whereas fibroblast-associated genes were used to define the reference profiles for the fibrosis-associated analysis. RCTD was then used to estimate the contribution of each reference profile to individual virtual spatial spots overlaid on the H&E-stained sections. The resulting estimates were used to identify and spatially map virtual spots enriched for SD1-associated or fibrosis-associated transcriptional programs. These enriched spatial distributions are referred to here as SD1-associated and fibrosis-associated virtual cell clusters using “doublet” mode, respectively.

### Single-cell gene expression data preprocessing, gene quantification, single-cell clustering and visualization

The single-cell raw sequencing reads were mapped against the human reference genome (GRCh38) using the Cell Ranger software (version 6.1.1). The filtered raw count matrix obtained from Cell Ranger was further processed using the Seurat (version 4.0.4) package^96^ in R. Cells with nFeature_RNA < 200, nFeature_RNA > 6000, nCount_RNA < 1000, min.cells < 3, min.features < 200, and percentage of mitochondrial genes > 15% were excluded in the following analysis. The filtered raw count matrices were normalized using the “NormalizeData” function. All samples were then integrated into a common subspace using canonical correlation analysis (CCA) based on the reciprocal PCA approach with default parameters. Total immune cells were then clustered using the “FindNeighbors” and “FindClusters” functions with a resolution of 0.4. After that, the clustered cells were projected into a two-dimensional space for visualization utilizing the “RunUMAP” function. The differentially expressed markers for each cluster were identified using the “FindMarkers” function with the ‘MAST’ algorithm. Cell clusters were manually annotated using canonical markers.

### Spatial transcriptome data preprocessing and gene quantification

Alignment and demultiplexing of raw sequencing reads were conducted via the Space Ranger pipeline, and subsequent analyses were conducted via Scanpy (version 1.10.4) in Python. We obtained the expression abundance information of different genes in different spots based on the genome comparison results, UMI counts and barcode sequence information.

### Spatial domain identification

MNMST^49^ (version 1.1) was utilized to identify spatial domains in spatial transcriptomics data by integrating gene expression and spatial location information of cells. In brief, MNMST jointly decomposes multi-layer networks to learn discriminative features of cells and identifies spatial domains by exploiting topological structure of affinity graph of cells.

### Principal Component Analysis (PCA) analysis of pseudo-bulk samples

Pseudo-bulk samples were created using averaged gene expression across all cells within this sample. PCA was then performed on these pseudo-bulk samples using “prcomp” function in R. The top 100 genes along PC1 axis were selected to explain the molecular mechanism underlying the sample distribution in PCA space.

### DEGs identification and gene enrichment analysis

The DEGs between the two groups were identified using the “FindMarkers” function with the “MAST” algorithm. Genes were defined as significantly upregulated if the average logarithm two-fold change (log2FC) > 0.58 and adjusted *P-value* < 0.05. The genes with log2FC <– 0.58 and adjusted *P*-value < 0.05 were considered significantly downregulated. The Gene Ontology (GO) enrichment analysis was performed using the “enrichGO” function in clusterProfiler^106^ package. Only the GO term of Biological Process (BP) was displayed.

### Single cell subpopulation re-integration

Innate T cells, CD8^+^ T cells, CD4^+^ T cells, B cells, myeloid cells, and epithelial cells were re-integrated and re-clustered using Seurat (version 4.0.4). The markers used for extracting these subpopulations were listed in Figure S1. Raw count matrices for each sample containing these subsets were extracted, re-integrated and re-clustered using the above procedure, separately. The parameter resolution was set to 0.6 when running the “FindClusters” function for analyzing these subsets.

### Evaluation of the transcriptional heterogeneity of single cell subpopulations

ROGUE^99^ is an entropy-based statistic to accurately quantify the purity of cell clusters. The transcriptional heterogeneity of each cell subpopulation was calculated as 1-ROGUE.

### Single-cell TCR and single-cell BCR analysis

The amino acid and nucleotide sequences of the TCR chains were assembled and annotated using the “cellranger vdj” function in Cell Ranger (version 6.1.1). Only cells with paired TCRα and TCRβ chains were included in the clonotype analysis. For cells with multiple VDJ annotations, we preserved only the VDJ genes with the highest UMI value. Cells sharing the same TCRα– and TCRβ-CDR3 amino acid sequences were assigned to the same TCR clonotype. The amino acid and nucleotide sequence of BCR chains were assembled and annotated by SCIGA^100^. The V gene somatic mutation rate (SHM) compared to the germline genes was also calculated using this software. Only cells with at least one productive heavy chain (IGH) and one productive light chain (IGK or IGL) were kept for further analysis. Cells sharing the same V/J gene and the same IGH and IGK/IGL-CDR3 amino acid were defined as a clonotype. The TCR clonotypes and the BCR clonotypes were integrated into transcriptome objects using barcode information. Shannon index (TCR/BCR diversity) of each sample was calculated using the “diversity” function in vegan package (version 2.5.6) in R.

### Pseudo-time trajectory inference

The undirected PAGA graph was generated utilizing a cluster-graph representation to capture the underlying topology between single cell clusters^65^. We performed pseudotime analysis using Monocle3^66^ (version 1.3.1). Briefly, Seurat object annotated with B cell clusters were fed directly into Monocle3, followed by the removal of batch effect using the “align_cds” function. Cell trajectory was calculated by using “learn_graph” function. Cell differentiation trajectory was inferred after dimension reduction and cell ordering with the default parameters. Gene expression along pseudotime data was extracted from the output of “plot_genes_in_pseudotime” function and was used to plot genes along pseudotime of lineages using ggplot2 (version 3.5.1) in the R package. The pseudotime values calculated by Monocle3 and the differentiation partition information are mapped to the original UMAP graph in Seurat.

### Lineage tree reconstruction from BCR clones

The Change-O toolkit (version 1.3.3)^103^ was utilized to contruct the BCR clonal trees. In brief, the C, V, D, J gene and CDR3 sequence for each BCR with assembled IgH sequence was firstly annotated by AssignGenes.py script. Only productive sequences were kept for downstream analyses. In each patient, the BCR sequences were clustered into clonal groups after determined the clustering threshold via a gamma/gamma mixture model of the nearest neighbor distances by the “findThreshold” function in the R package SHazaM (version 1.0.2). The DefineClones.py script was then applied to group similar IgH sequences into clonal groups based on IGHV alleles, IGHJ alleles, junction length and junction distance. Before that the threshold was calculated by SHazaM in each patient. Germline sequences were reconstructed by CreateGermlines.py script. For the selected representative BCR clones, we first used the “msaConsensusSequence” function from the R package msa^104^ (version 1.34.0) to obtain a consensus CDR3 nucleotide acid sequence for each clonal group as the root sequence in the tree building. Next, lineage trees were constructed via maximum parsimony employing the “buildPhylipLineage” function from the R package Alakazam (version 1.0.2). The cell type composition of each node was depicted by a pie chart.

### Scoring cells using gene expression signatures

The “AddModuleScore” function of Seurat was applied with default parameters to score each signature (Table S5) in each T cell^112^, B cell^58^, and myeloid cell population^113^ and the mean score of all cells in each subset was calculated.

### Cell-cell communication analysis

The cell-cell interactions from scRNA-seq data were inferred and visualized by CellChat^77^ (version 2.1.2), which models the probability of cell-cell communication by integrating gene expression with prior knowledge of the interactions between signaling ligands, receptors, and their cofactors. The interactions in datasets “Secreted Signaling” were shown in this study. Cell-cell communications from spatial transcriptomics data were inferred by COMMOT^98^ (version 0.0.3), which screens spatial cell-cell talks using collective optimal transport method to handle complex molecular interactions and spatial constraints. The “commot.tl.spatial_communication” function was utilized construct cell-cell communication networks for the ligand-receptor pairs with a spatial distance constraint of 500 μm. Niche interactions and communication analysis of interest were performed using NICHES software^107^.

### Ro/e index calculation

The group distribution preference of scRNA-seq and snRNA-seq cell cluster was indicated by Ro/e index.

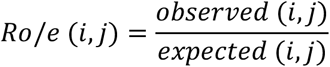

Where *observed* (*i*, *j*) represents the cell number of a given subpopulation *i* in a specific disease group *j*. The *expected* (*i*, *j*) is calculated using the following formula:

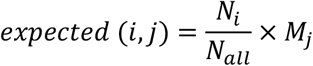

*N_i_* is the cell number of the given subpopulation *i*. *M_j_* is the cell number of all subpopulations of a given group *j*. *N_all_* is the total cell number of the selected subpopulation. An Ro/e value greater than one indicates that a subpopulation in a specific group is more frequently observed than random expectations.

### Cell population relative enrichment score calculation and spot deconvolution

For each cell subpopulation *i* in scRNA-seq data, we calculated their relative enrichment score in spatial domain *j* using the cell markers as listed in Table S4.

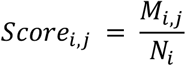

*N_i_* is the total number of spots with UMI > 1 of all the markers in subpopulation *i*. *M_i,j_* is the number of spots in spatial domain *j* with UMI > 1 of all the markers in subpopulation *i*. Cell types from spatial mixtures were also decomposed by STRIDE^50^ (version 0.0.2), leveraging latent dirichlet allocation, a generative probabilistic model, for the discovery of functional topics from snRNA-seq data.

### Virtual spots clustering

To delineate spatial functional domains (e.g., TLS), we performed unsupervised graph-based clustering on the predicted gene expression matrix of all virtual spots. Dimensionality reduction, Shared Nearest Neighbor (SNN) graph construction, and spatial clusters were performed using the above procedure. Spatial clusters were color-coded and projected back onto the high-resolution H&E images for histological validation.

## Statistical analysis

Statistical analysis was performed using the R 4.3.3 framework and GraphPad Prism software. Paired or unpaired Student’s t-test, Wilcoxon’s sign rank test, and Wilcoxon’s rank-sum test were applied as appropriate for the data analysis. Adjusted *p* values were calculated by the Benjamini Hochberg method to control the false discovery rate. The significance levels of the performed tests were denoted by asterisks: \*\*\*\**P* < 0.0001, ***P < 0.001, \*\**P* < 0.01, and \**P* < 0.05.

## Declaration of generative AI and AI-assisted technologies in the manuscript preparation process

During the preparation of this work, the authors used ChatGPT for minor rephrasing of the text. After using this tool, the authors reviewed and edited the content as needed and take full responsibility for the content of the publication.

## Supplementary Figures

**Figure S1.**
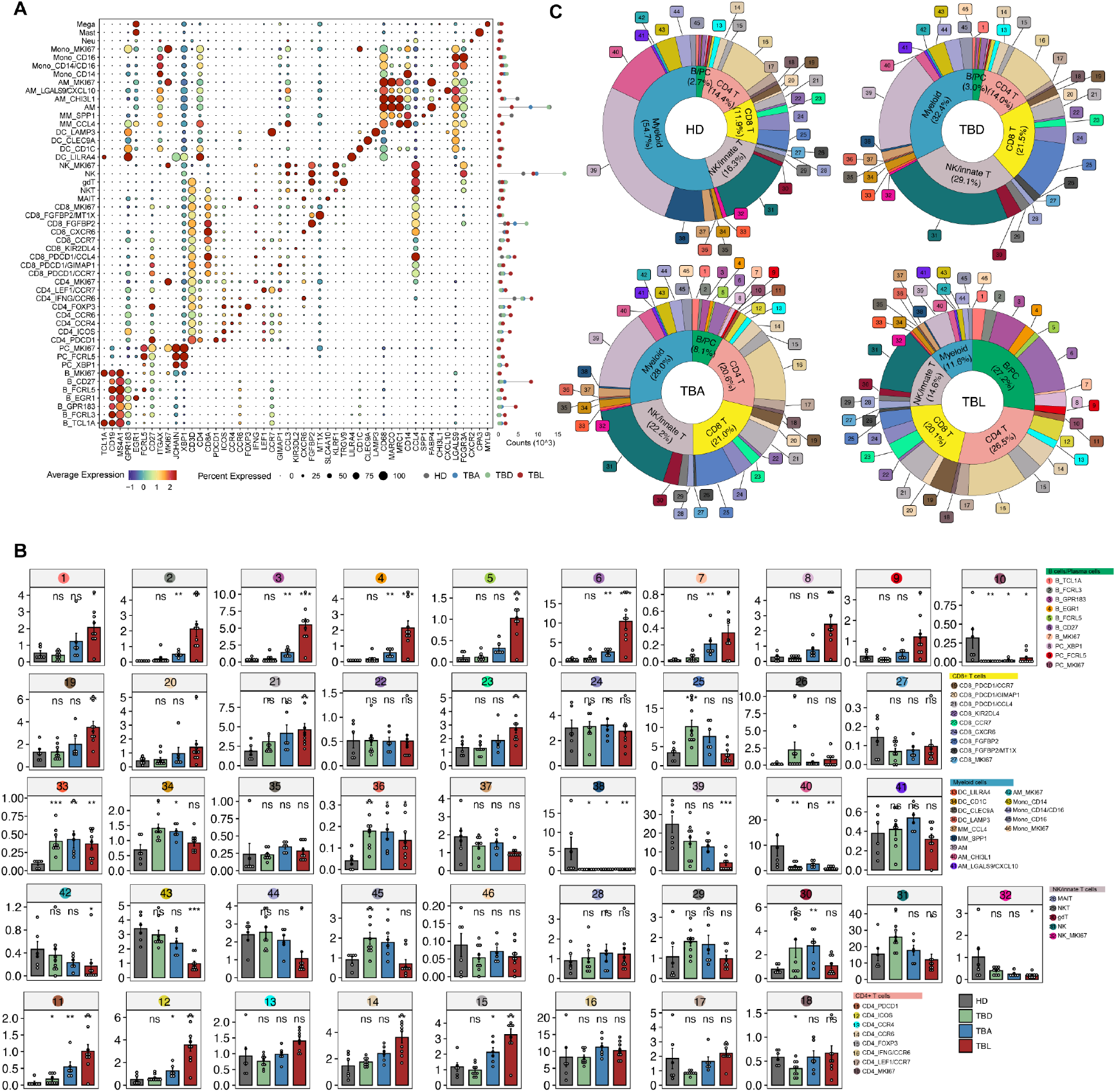
Characteristics of scRNA-seq cell subsets, related to Figure 1. (A) Canonical markers used to define the 49 immune cell subtypes. (B) Fraction of each cell subtype (Mann-Whitney U Test, \**P* < 0.05, \*\**P* < 0.01, \*\*\**P* < 0.001). (C) Pie charts showing the relative cellular composition of HD, TBD, TBA and TBL samples. HD, healthy donors; TBD, tissues distal to the lesions; TBA, tissues from lesion boundaries or adjacent areas; TBL, active lesion tissue.

**Figure S2.**
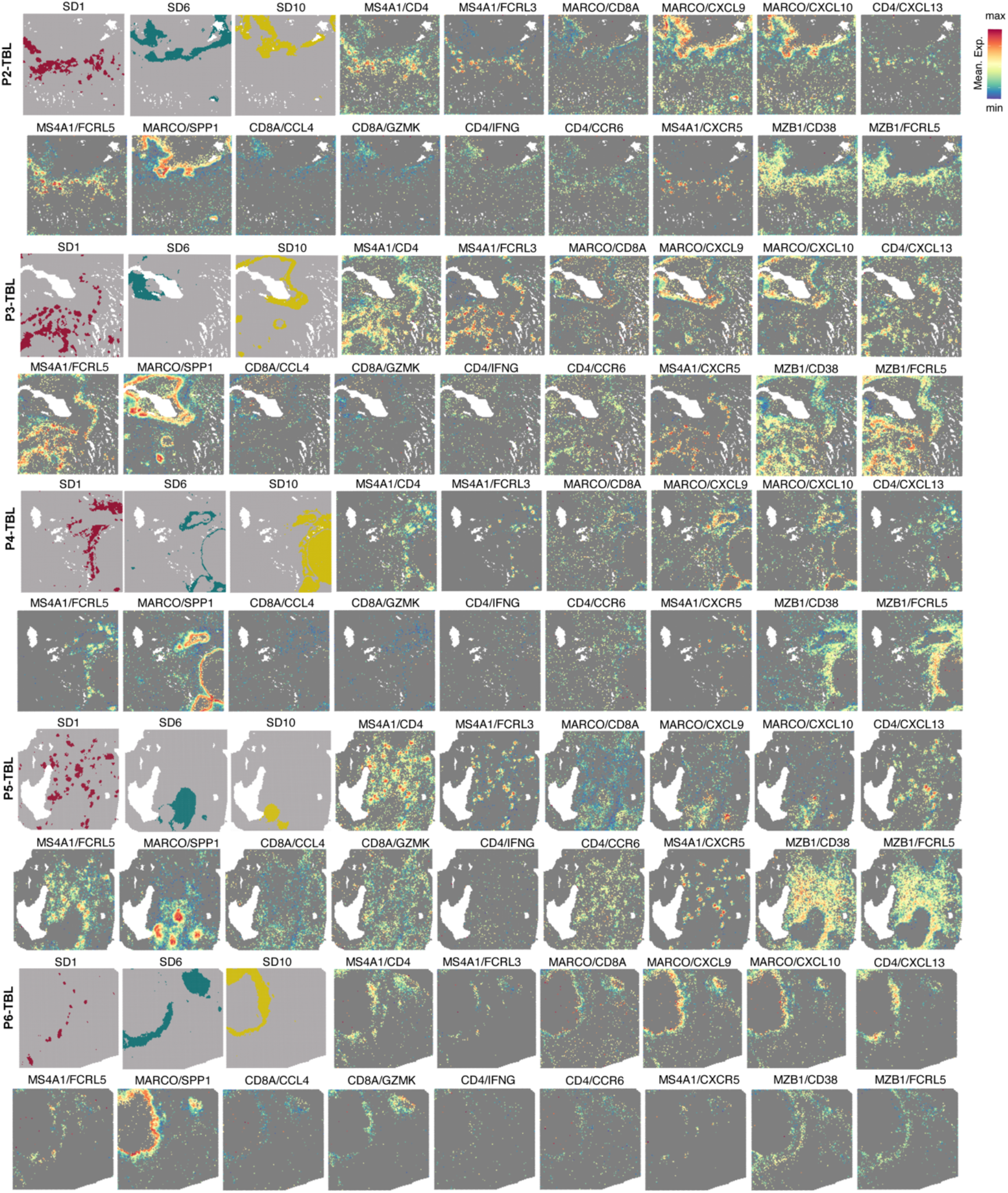
Spatial expression of indicated genes across granulomas, related to Figure 2. Spatial heatmaps showing the mean expression of each two-marker combination in P2-TBL to P6-TBL samples.

**Figure S3.**
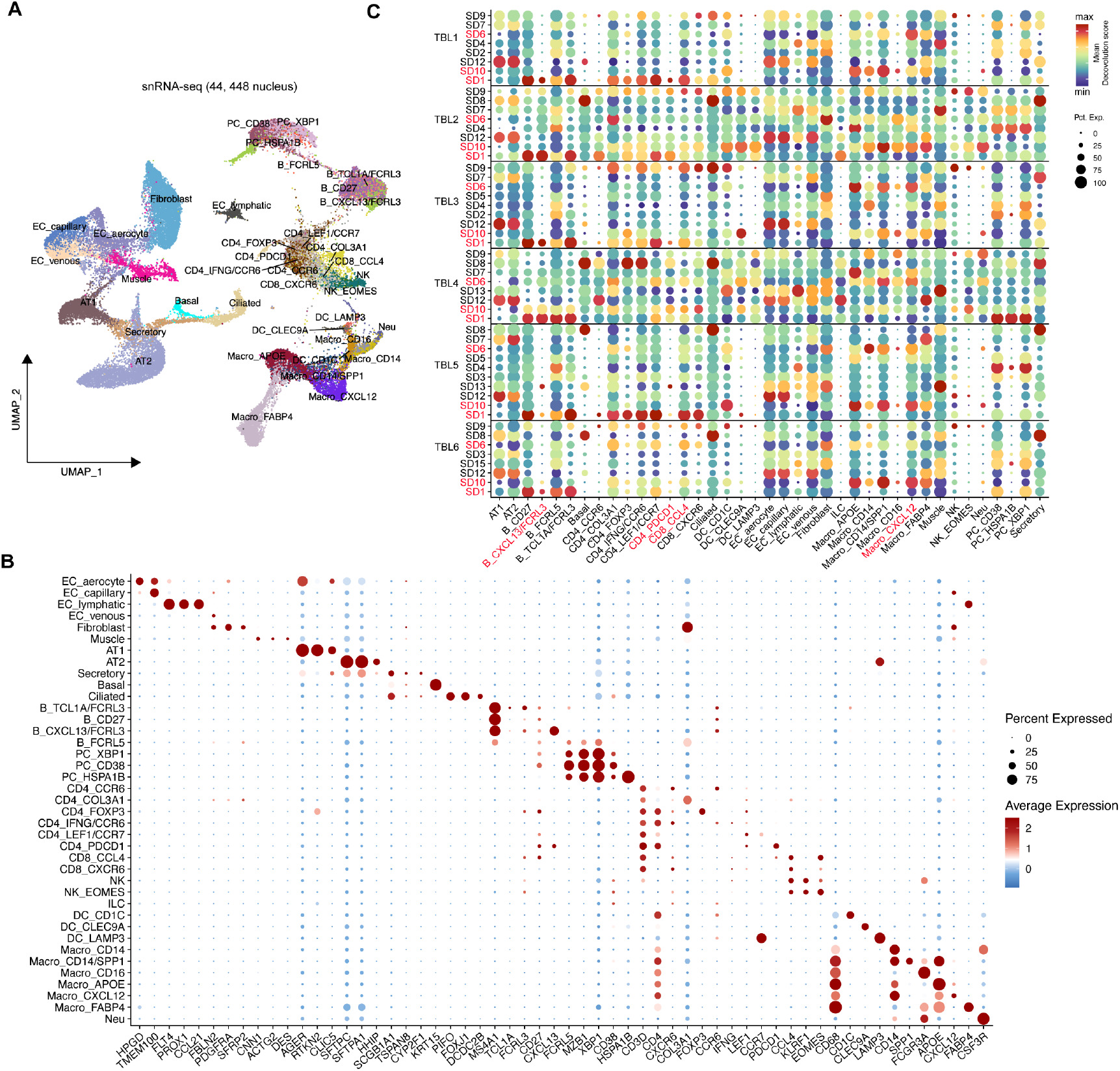
Characteristics of snRNA-seq cell subsets, related to Figure 2. (A) The cell subtypes identified by snRNA-seq. A total of 39 cell subtypes were resolved. (B) Canonical markers used to define the 39 cell subtypes. (C) Deconvolution scores for all cell types mapped to spatial locations.

**Figure S4.**
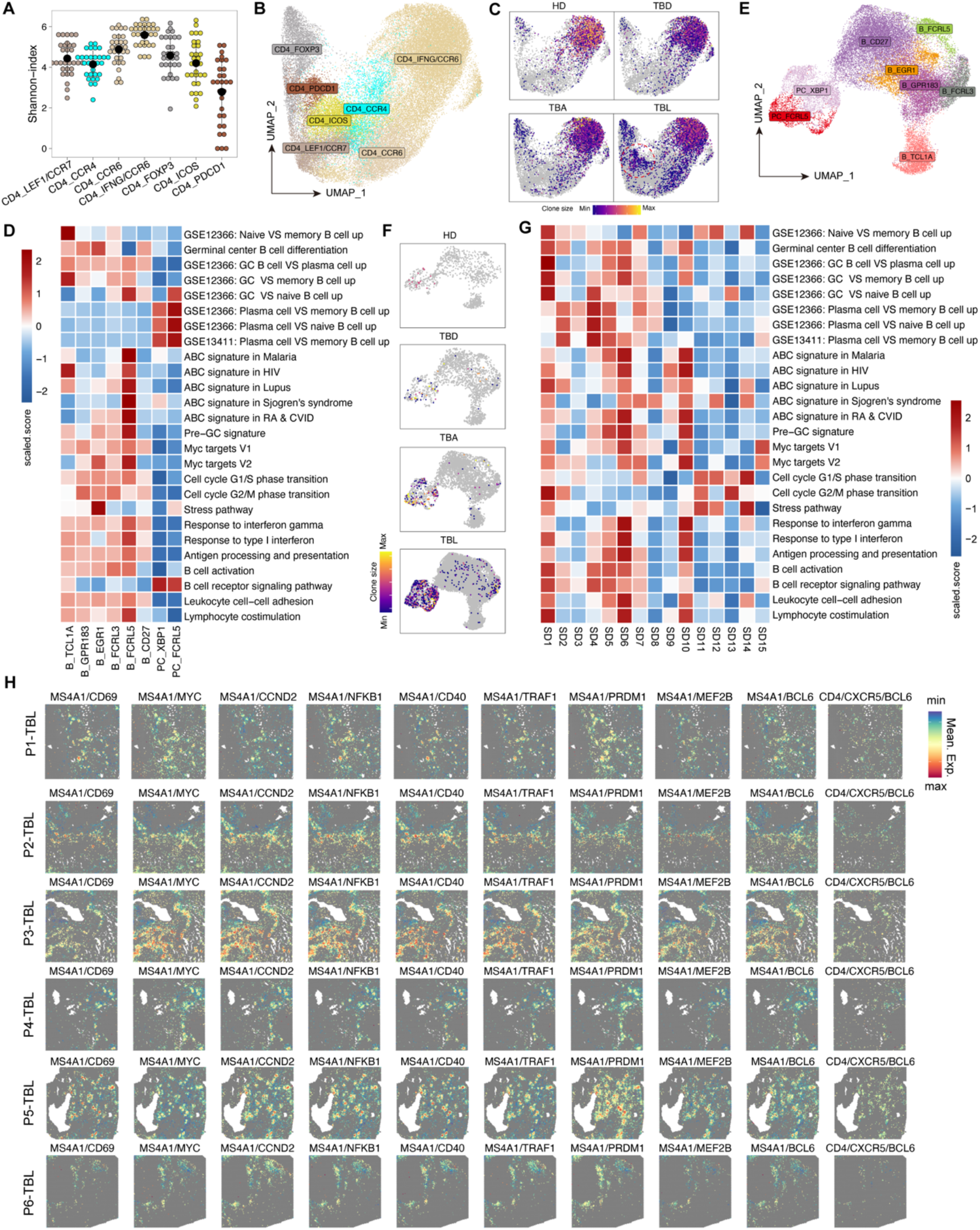
T_FH_-B cell interplay niches and their functional signatures, related to Figure 3. (A) TCR diversity, measured using the Shannon index, across CD4^+^ T-cell subsets. (B) UMAP projection of CD4^+^ T-cell clusters. (C) UMAP showing TCR clone size distributions in CD4^+^ T cells from HD, TBD, TBA, and TBL samples. (D) Heatmap showing gene-set-based phenotypic scores across B-cell subsets. (E) UMAP projection of B-cell clusters. (F) UMAP showing BCR clone size distributions in B cells from HD, TBD, TBA, and TBL samples. (G) Heatmap showing B-cell-related gene-set scores across spatial domains. (H) Spatial heatmaps showing the mean expression of GC-related marker pairs. HD, healthy donors; TBD, tissues distal to the lesions; TBA, tissues from lesion boundaries or adjacent areas; TBL, active lesion tissue.

**Figure S5.**
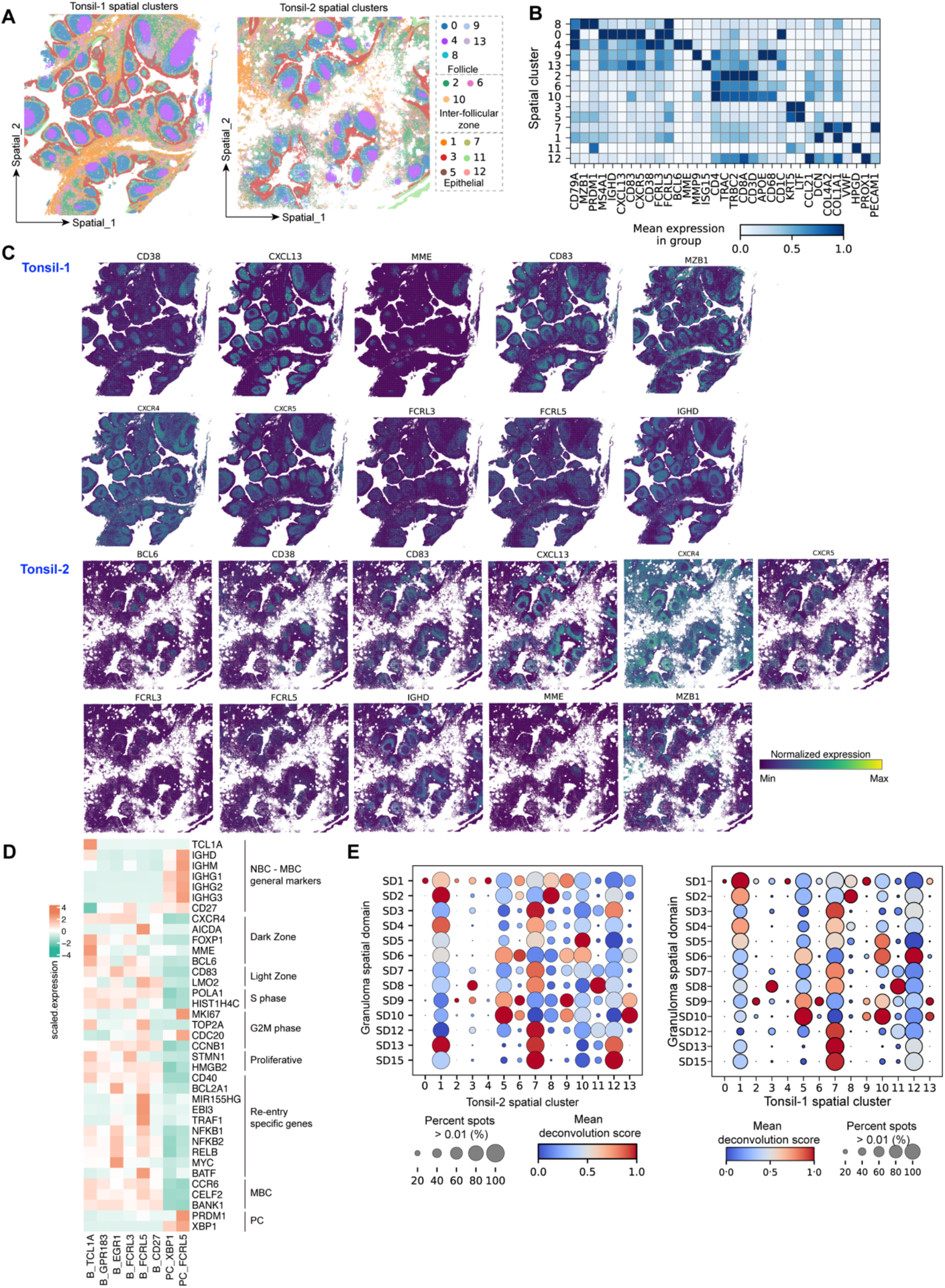
Spatial transcriptomic architecture of tonsil tissue. (A) Spatial clusters identified in two normal tonsil sections. (B) Heatmap showing representative marker expression in each spatial cluster. (C) Spatial heatmaps showing representative marker expression patterns across tissue sections. (D) Mean expression of GC-related genes across B-cell subsets. (E) Mean deconvolution score for each tonsil spatial cluster projected onto spatial domains in human granulomas.

**Figure S6.**
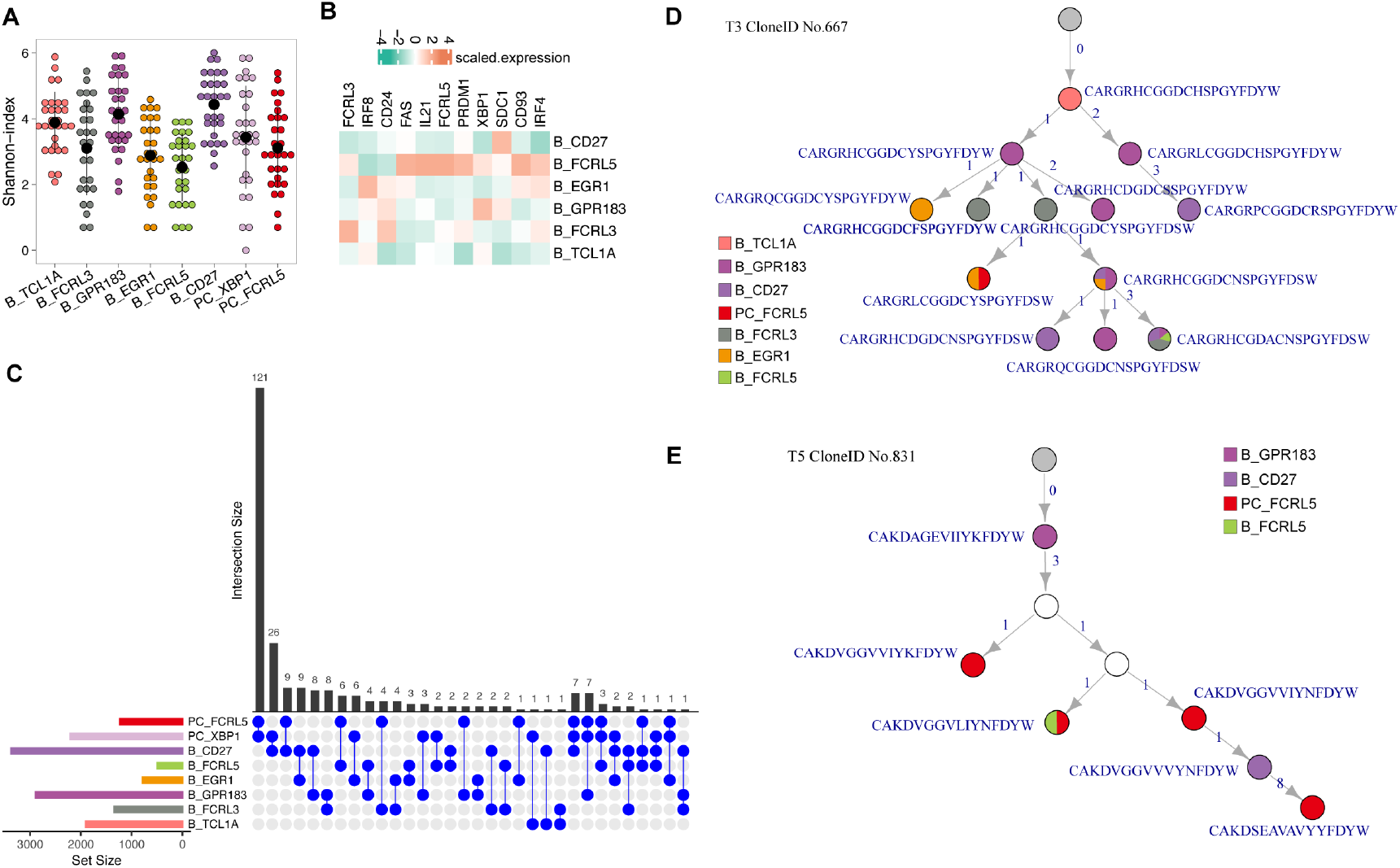
BCR-based differentiation trajectory of PC_FCRL5 cells within TLS. (A) BCR diversity, measured by the Shannon index, across B-cell subsets. (B) Heatmap showing the mean expression of PC differentiation-associated genes across B-cell subsets. (C) Upset plot showing shared BCR clones among B-cell subsets. (D-E) Representative BCR lineage trees from patients T3 (D) and T5 (E), annotated with CDR3 amino acid sequences, cell counts on the edges, and subtype composition shown as pie charts.

**Figure S7.**
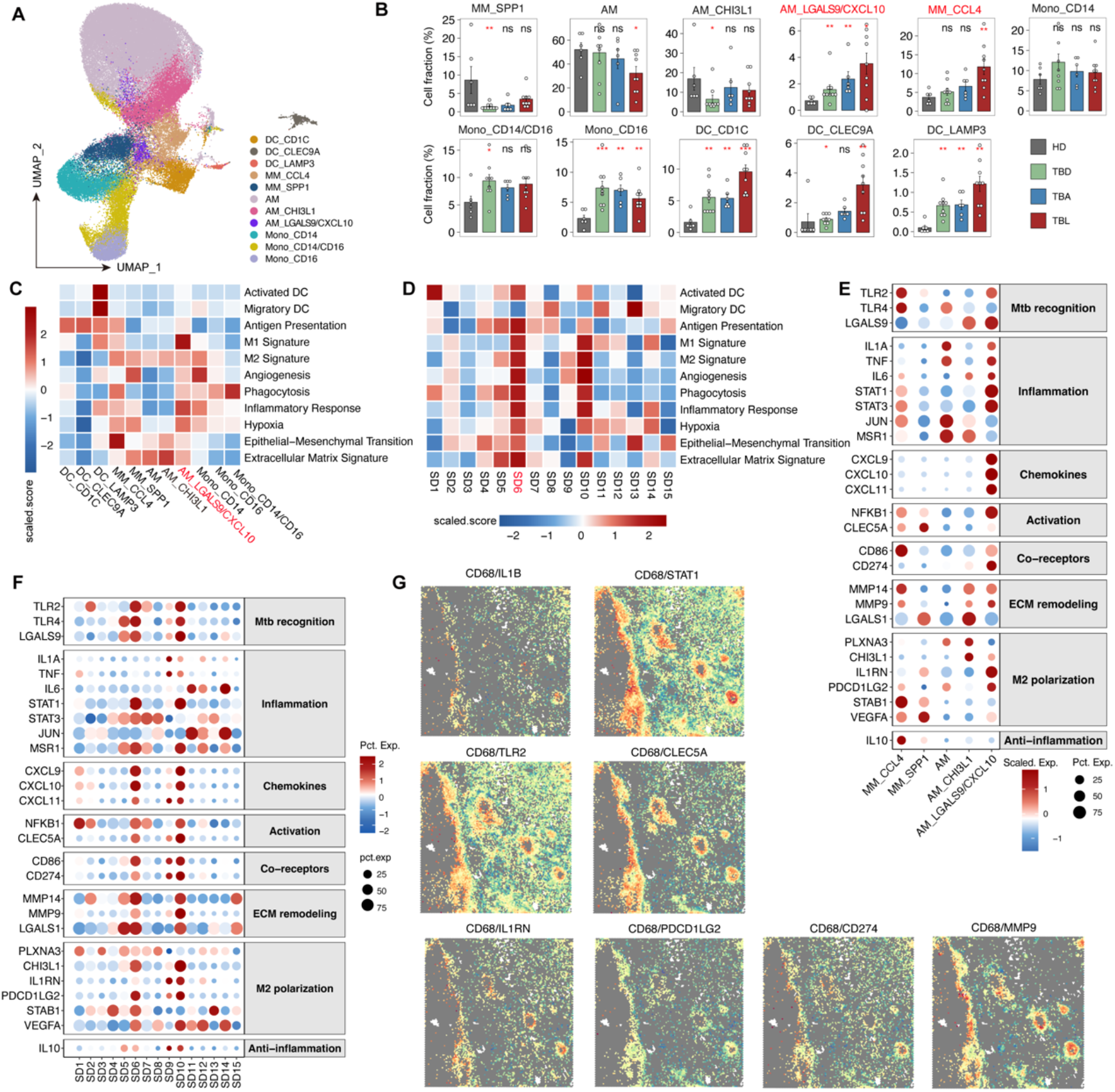
Transcriptional phenotypes of macrophage populations in TB granulomas. (A) UMAP projection of myeloid-cell clusters. (B) Fractions of myeloid-cell populations. (C) Heatmap showing gene-set-based molecular phenotypes across myeloid subtypes. (D) Heatmap showing functional gene-set scores for myeloid cells across spatial domains. (E-F) Dot plots showing the expression of selected functional genes across macrophage populations (E) and spatial domains (F). (G) Spatial heatmaps showing the mean expression of selected marker combinations in P1-TBL.

**Figure S8.**
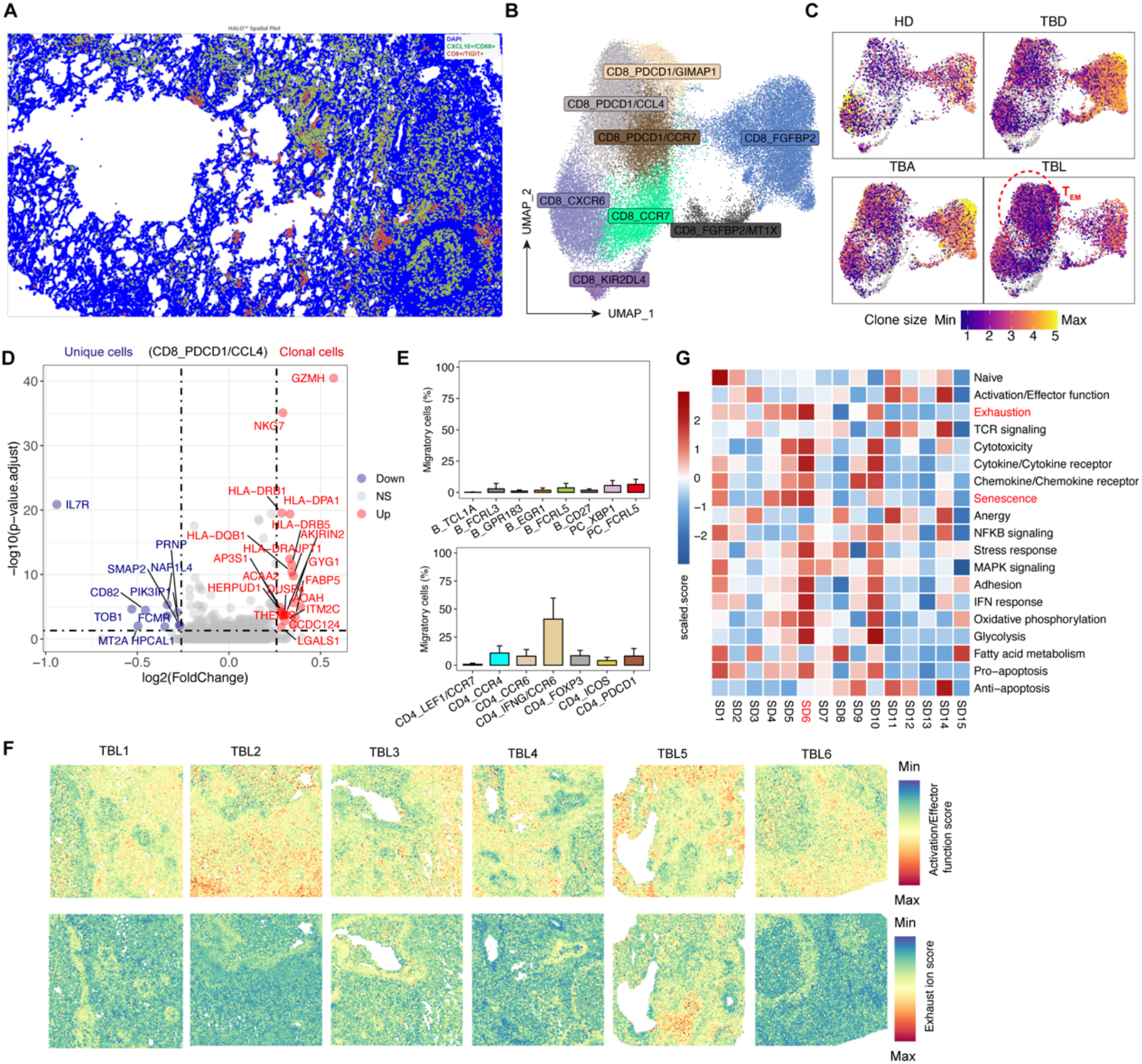
Transcriptional signatures of CD8^+^ T-cell subsets in TB granulomas. (A) Histocytometry of lung sections from patients with TB showing CD8^+^TIGIT^+^ (red) and CXCL10^+^CD68^+^ macrophages (green). (B) UMAP projection of CD8^+^ T-cell clusters. (C) UMAP showing TCR clone size distributions in CD8^+^ T cells from HD, TBD, TBA, and TBL samples. (D) Volcano plot showing differentially expressed genes between clonal and non-clonal T_EM_ cells. (E) Bar plot showing the fraction of migratory cells across CD4^+^ T-cell and B-cell subtypes. (F) Spatial heatmaps showing exhaustion and activation/effector scores in the six TB patients. (G) Heatmap showing CD8^+^ T cell functional gene-set scores across spatial domains. HD, healthy donors; TBD, tissues distal to the lesions; TBA, tissues from lesion boundaries or adjacent areas; TBL, active lesion tissue.

**Figure S9.**
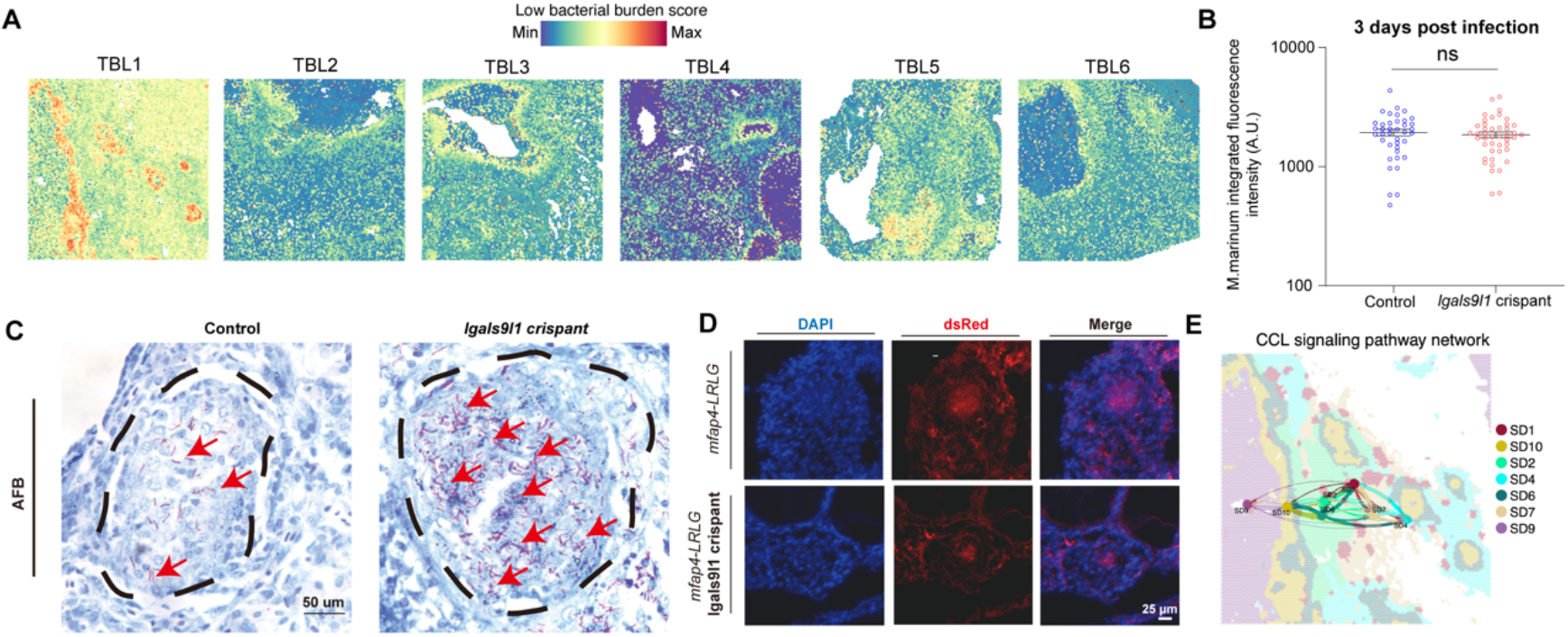
Function of LGALS9^high^ chemokine-expressing macrophages, related to. Figure 5. (A) Spatial heatmaps showing low bacterial burden scores in six patients. (B) Comparison of bacterial loads at 2 hours and 3 days post-infection between AB larvae and crispant larvae whose *lgals9l1* gene were knocked out at the fertilized embryo stage. Each point represents the bacterial fluorescence intensity in an individual zebrafish larva. (C) Representative images of AFBs in granulomas from the Control and *lgals9l1* crispant groups. Black dashed lines outline granulomas, and red arrows indicate AFB signals. (D) Distribution of macrophages in granulomas from the Control and *lgals9l1* crispant groups. (E) Spatial communications between domains inferred by COMMOT in P1-TBL.

**Figure S10.**
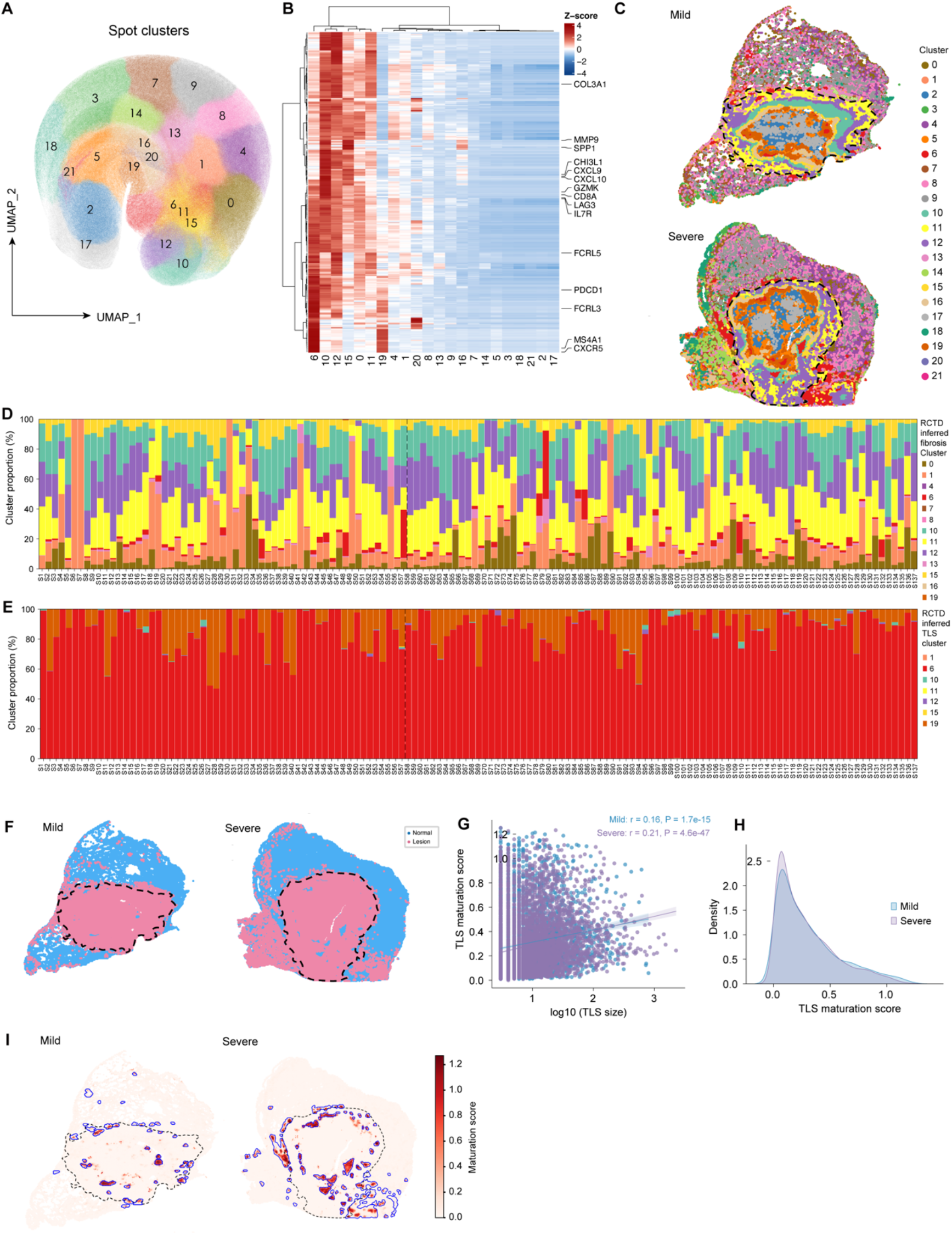
Virtual-cell clustering and functional annotation, related to Figure 6. (A) UMAP projection of virtual-cell clusters across 137 TB samples. (B) Heatmap showing representative marker expression across virtual-cell clusters. (C) Spatial projection of virtual-cell clusters onto sections from representative mild and severe TB samples. Dashed lines indicate inferred fibrotic boundaries. (D) Stacked bar plot showing the relative fractions of inferred fibrosis clusters in each TB sample, estimated using RCTD with the fibroblast cluster from the snRNA-seq data as the reference. (E) Stacked bar plot showing the relative fractions of inferred TLS clusters in each TB sample, estimated using RCTD with SD1 spatial transcriptomic signatures as the reference. (F) Spatial distribution of normal and lesion regions in sections from representative mild and severe TB samples. Dashed lines indicate inferred fibrotic boundaries. (G) Pearson correlation between TLS size and TLS maturation score (defined by average expression of *CR2* and *FCER2*) in mild and severe TB samples. Each point represents one TLS structure. (H) Density plots showing the distributions of TLS maturation scores in mild and severe TB samples. (I) Spatial distribution of TLS maturation scores on representative mild and severe TB sections.

